# Framework estimation of stochastic gene activation using transcription average level

**DOI:** 10.1101/2021.09.01.458497

**Authors:** Liang Chen, Genghong Lin, Feng Jiao

**Author notes:** Corresponding author: F.J. L.C. and G.L. contributed equally.

## Abstract

Gene activation is usually a non-Markovian process that has been modeled as various frameworks that consist of multiple rate-limiting steps. Understanding the exact activation framework for a gene of interest is a central problem for single-cell studies. In this paper, we focus on the dynamical data of the average transcription level *M* (*t*), which is typically neglected when deciphering gene activation. Firstly, the smooth trend lines of *M* (*t*) data present rich, visually dynamic features. Secondly, tractable analysis of *M* (*t*) allows the establishment of bijections between *M* (*t*) dynamics and system parameter regions. Because of these two clear advantages, we can rule out frameworks that fail to capture *M* (*t*) features and we can further test potential competent frameworks by fitting *M* (*t*) data. We implemented this procedure to determine an exact activation framework for a large number of mouse fibroblast genes under tumor necrosis factor induction; the cross-talk between the signaling and basal pathways is crucial to trigger the first peak of *M* (*t*), while the following damped gentle *M* (*t*) oscillation is regulated by the multi-step basal pathway. Moreover, the fitted parameters for the mouse genes tested revealed two distinct regulation scenarios for transcription dynamics. Taken together, we were able to develop an efficient procedure for using traditional *M* (*t*) data to estimate the gene activation frameworks and system parameters. This procedure, together with sophisticated single-cell transcription data, may facilitate a more accurate understanding of stochastic gene activation.

**Author Summary:** It has been suggested that genes randomly transit between inactive and active states, with mRNA produced only when a gene is active. The gene activation process has been modeled as a framework of multiple rate-limiting steps listed sequentially, parallel, or in combination. The system step numbers and parameters can be predicted by computationally fitting sophisticated single-cell transcription data. However, current algorithms require a prior hypothetical framework of gene activation. We found that the prior estimation of the framework can be achieved using the traditional dynamical data of mRNA average level *M* (*t*) which present easily discriminated dynamical features. The theory regarding *M* (*t*) profiles allows us to confidently rule out other frameworks and to determine optimal frameworks by fitting *M* (*t*) data. We successfully applied this procedure to a large number of mouse fibroblast genes and confirmed that *M* (*t*) is capable of providing a reliable estimation of gene activation frameworks and system parameters.

## 1 Introduction

Gene transcription is a random process in virtually all genomic loci, for which messenger RNA (mRNA) molecules for active genes are produced in a bursting fashion in which an episode of transcriptional activity is interrupted by irregular gene inactivation periods [1–3]. A central problem in the study of stochastic gene transcription has been understanding regulation scenarios that control random gene activation (*on*) and inactivation (*off*) in response to environmental changes [4–7]. The tremendous effort expended in this endeavor over the last few decades has generated massive amounts of data at the single-cell level and has produced many important observations [8–10].

Real-time imaging of transcriptional bursting makes it possible to count the durations of each gene *on* and *off* period along the entire timeline, which generates duration distributions for both gene *on* and *off* periods, respectively. For instances of the *Escherichia coli* P_lac*/*ara_ promoter [11] and yeast *FLO11* genes [12], their *on* and *off* periods are all well fitted by single exponential distributions. These observations support the classical two-state model shown in Fig. 1a, that the gene for turning genes *on* and *off* are all controlled by single rate-limiting biochemical steps, with synthesis of mRNA when the gene is *on* and degradation of the gene all being controlled by single rate-limiting steps [1, 3]. The exponentially distributed *on* period is one of the few universal features of transcription present in both prokaryotic and eukaryotic genes [6, 8, 13]. However, the duration of the *off* state is highly gene-specific, and is manifested by the observed gamma distribution with a unique peak for mouse fibroblast genes [8] and *E. coli tetA* promoters [14]. Gamma distribution or more general non-Markovian processes can be mathematically explained by assuming that the gene for activating the *on* period is directed by a single pathway consisting of multiple sequential rate-limiting steps (e.g., the three-state model in Fig. 1b) [15], the parallel competitive rate-limiting pathways (e.g., the cross-talking pathways model in Fig. 1c) [16], or a combination of both (e.g., Fig. 1d) [17]. However, these theoretical approaches are unable to determine the best framework to describe the observed non-Markovian gene activation, although they have provided good approximations to the downstream distribution of gene transcription [18, 19].

**Figure 1:**
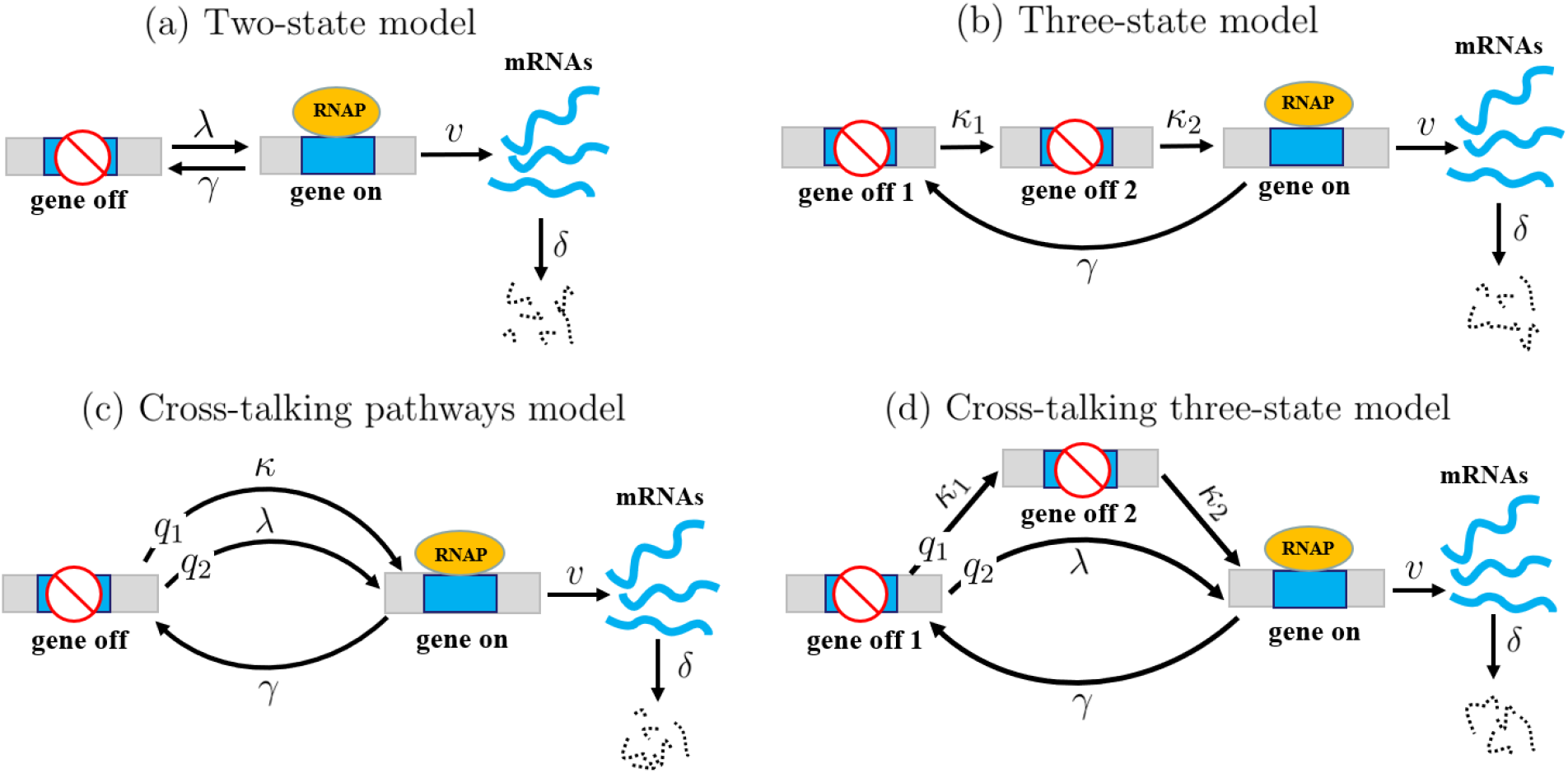
Different frameworks that direct stochastic gene activation (*on*). Other processes of gene inactivation (*off*), mRNA synthesis when the gene is *on*, and mRNA degradation are all determined by single rate-limiting steps at constant rates. (a) Two-state model. The gene is activated through a single rate-limiting step at a constant rate. (b) Three-state model. Gene activation is regulated by two sequential rate-limiting steps at a constant rate. (c) Cross-talking pathway model. The gene is activated by two different competitive rate-limiting pathways with selection probabilities *q*_1_ and *q*_2_ of the two pathways satisfying *q*_1_ + *q*_2_ = 1. (d) Cross-talking three-state model. The gene is activated either by a pathway consisting of two sequential rate-limiting steps, or alternatively, by a single rate-limiting pathway, with constant rates.

The snapshot data for the distribution histogram of mRNA copy numbers in an isogenic cell population at different time points carry rich dynamic information on fluctuations in transcription [20]. When combined with mathematical models, the fit of mRNA (or other RNA types) distribution data has served as a powerful tool for revealing the multi-step regulation of the activation of different genes in bacteria, yeast, and human cells [9, 20, 21]. However, the calculation of exact forms of dynamical mRNA distribution requires solving infinite arrays of chemical master equations under the whole parameter region of the models, which is beyond the scope of standard theoretical methods, even for the simplest two-state model (Fig. 1a) [22–24]. Fitting of mRNA distribution data must integrate several computational tools to determine the rate-limiting step numbers in the activation framework and search for suitable system parameters [9, 20]. However, current computation algorithms focus only on a class of prior hypothetical multi-step gene activation; therefore, additional competent frameworks may be ignored. Moreover, some typical dynamical transition patterns among the different mRNA distribution modes can be well exhibited by all the models in Fig. 1 [24–26], preventing a direct way to rule out models that do not panoramically match the transition patterns of dynamical mRNA distribution.

The steady-state measurement of gene transcription under different cellular conditions has generated a large dataset of mRNA distribution and its mean level *M* , the Fano factor *ϕ* (the variance over *M*), and noise *CV* ^2^ (*ϕ* over *M*) [1,10]. By virtue of mathematical models, fitting steady-state data has revealed a large spectrum of regulation scenarios that cells utilize in response to environmental changes [3,10]. The steady-state mRNA distributions observed so far are often shown to be decaying, unimodal, or bimodal [1, 3]. However, the models in Fig. 1 can only generate the three distribution modes shown at steady-state [15, 23, 27, 28], suggesting that the limited mRNA distribution modalities are insufficient to map reversely onto the diversified frameworks of gene activation. The steady-state data of noise *CV*^2^, Fano factor *ϕ*, and mean level *M* , when mapped as scattered points onto *M*-*CV*^2^ and *M*-*ϕ* planes, provide a diagram of trend lines of *CV*^2^ and *ϕ* against *M* under varying environments [2,10]. For a given gene of interest in *E. coli*, yeast, or mammalian cells, the trend lines fitted by different models have revealed distinct regulation scenarios [29]. However, the scenario that plays a dominant role in gene regulation remains elusive.

In contrast to the time-consuming single-cell measurements that require RNA labeling and imaging with high sensitivity and resolution [11, 21], the dynamical mRNA average level *M* (*t*) can be relatively easily captured by conventional methods at the cell population level [21,30]. Previous studies have revealed rich temporal profiles of *M* (*t*) for different genes and cellular conditions, such as monotonic increases in the *E. coli* promoter P_lac*/*ara_ [11], up-and-down behavior in the *c-Fos* gene in human osteosarcoma [21], multiple peaks in mouse fibroblast genes [30, 31], and even oscillations in yeast stress-induced genes [32]. These observations give rise to the problem of whether such rich dynamical behaviors of *M* (*t*) can be mapped back to the diversified frameworks of gene activation. To achieve this goal, the key objective is to establish bijections between the dynamical features of *M* (*t*) and the parameter regions for certain mathematical models. This allows us to rule out models that do not capture the exhibited dynamical features of *M* (*t*) and to test the simplest of the remaining models on the basis of their fit to *M* (*t*) data. In this study, we assumed that gene activation is regulated by a combination of sequential and parallel pathways, as shown in Fig. 1, and we illustrated the way that dynamic mRNA average level data can be utilized to help estimate the gene activation frameworks.

## 2 Results

### 2.1 Cross-talking three-state model

To make the paper easier to follow, we focused on analyzing the mRNA average level *M* (*t*) data from a large group of mouse fibroblast genes under cytokine tumor necrosis factor (TNF) stimulation conditions [30, 31]. With the exception of the simple monotonic growth of *M* (*t*) generated by late response genes, the rich non-monotonic behaviors of *M* (*t*) have also been determined, such as up-and-down for the *Fos* gene, up-down-up for the *Cxcl1* gene, and damped oscillation with multiple peaks for the *Nfkbia* gene.

To determine the framework that can best capture the rich transcription dynamics of mouse fibroblast genes, theoretical bijections between the dynamical features of *M* (*t*) and three mathematical models were established (Table 1). The two-state model shown in Fig. 1a can only generate monotonic increasing dynamics of *M* (*t*) [29], and thus is not suitable for the discussion of non-monotonic dynamical behaviors. The three-state model shown in Fig. 1b is proven to display damped oscillatory dynamics of *M* (*t*) under a certain parameter region [33, 34]. However, such oscillation behavior is almost invisible owing to its rapid exponential decay, and only slightly slows down the dynamic increase in *M* (*t*) [34]. The frameworks with two or more parallel pathways can capture the up-and-down dynamics of *M* (*t*), but fails to generate more complex transcription dynamics [29, 35]. Moreover, the cross-talking pathways model (Fig. 1c) generates up-and-down *M* (*t*) only when the stronger pathway is frequently selected to active the gene [29], which is incompatible with the robust up-and-down dynamics of *M* (*t*), even if the TNF induction level is extremely low [31]. Taken together, the activation frameworks of a single pathway or parallel pathways alone are insufficient to capture the rich dynamics of *M* (*t*) from mouse fibroblast genes (Table 1).

**Table 1:** Bijection theory between *M* (*t*) dynamical features and system parameter regions for the (a) two-state [29], (b) three-state [33, 34], (c) cross-talking pathways [29] and (d) multiple pathways models [35]. *α, ξ,* Λ, *α*_1_, and *x*_1_ are the auxiliary numbers associated with the system parameters, and *γ* and *δ* are the gene inactivation rate and mRNA degradation rate, respectively. *a*The three-state model generates a damped oscillatory *M* (*t*) when *α*^2^ < *ξ* [33, 34]. However, such oscillation decays exponentially and displays visually increasing dynamics [34].

| Mathematical models | $M(t)$ dynamical profiles $\iff$ | Parameter regions |
| --- | --- | --- |
| (a) Two-state model | Increase | All parameters |
| (b) Three-state model | Increase | $\alpha^2 \geq \xi$ |
| | Almost increase<sup>a</sup> | $\alpha^2 < \xi$ |
| (c) Cross-talking pathways model | Increase | $\Lambda \geq \min\{\delta, \gamma\}$ |
| | Up-and-down | $\Lambda < \min\{\delta, \gamma\}$ |
| (d) Multiple pathways model | Increase | $x_1 \geq \min\{\delta, \alpha_1\}$ |
| | Up-and-down | $x_1 < \min\{\delta, \alpha_1\}$ |

By combining the three-state model (Fig. 1b) and cross-talking pathways model (Fig. 1c), it is possible to generate new dynamic *M* (*t*) features. The simplest combination is shown in Fig. 1d, which we call the cross-talking three-state model. This model can be viewed as adding a parallel pathway in the three-state model or decomposing a pathway of the cross-talking pathways model into two sequential steps, as shown in the following scheme:

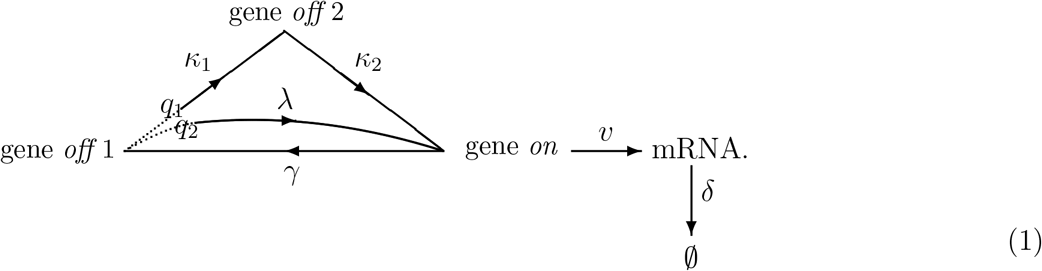

We assumed that the gene is activated by two competitive pathways. These are the weak basal pathway, which is has a selection probability *q*_1_ and consists of two sequential rate-limiting steps with strength rates *κ*_1_ and *κ*_2_, or alternatively the strong rate-limiting signaling pathway, which has a selection probability *q*_2_ and strength rate *λ*, satisfying:

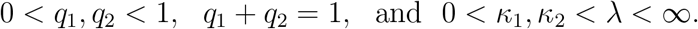

The basal pathway is regulated independently by a spontaneous mechanism to maintain basal transcription levels under normal cellular growth conditions [36, 37]. The assumption of two sequential steps and small strength rates for the basal pathway is in close agreement with the real-time imaging data of the *off* period of 16 mouse fibroblast genes [8]. Moreover, if *κ*_1_ or *κ*_2_ is relatively large, then the basal pathway can be mathematically approximated by a single rate-limiting step [33], and the framework (1) reduces to the cross-talking pathways model (Fig. 1c). A stronger signaling pathway is triggered when cells receive external cues, and downstream transcription factors (TFs) are activated by special signal transduction pathways to upregulate gene transcription [31, 38]. For each target gene, its activation is ultimately mediated through the binding of downstream TFs in the basal or signaling pathways at the cognate DNA sites in the gene promoter or enhancer domains [5, 38]. The selection probabilities *q*_1_ and *q*_2_ may then quantify the concentration and availability of activated TFs in each pathway to competitively form TF/DNA binding configurations, while the inducible activation rate *λ* of the signaling pathway quantifies the binding accessibility and strength between the corresponding TFs and DNA sites [5, 17, 35, 38].

### 2.2 Dynamics of *M* (*t*) and the bijection with parameter regions

To establish the bijection between the *M* (*t*) profiles and the parameter regions of the model (1), we first need to calculate the exact forms of *M* (*t*) in terms of system parameters. At time *t* ≥ 0, let random variable *X*(*t*) = *X* = *o*_11_, *o*_12_, *o*_2_, *e*, specify the states of gene *off* 1 for basal pathway, gene *off* 1 for signaling pathway, gene *off* 2, and gene *on*, respectively. Then define:

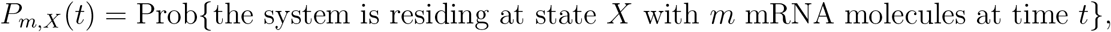

and the mass function:

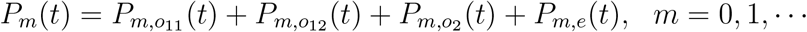

that quantifies the probability of *m* mRNA transcripts at time *t* in a single cell.

Following the standard procedure, we can obtain an infinite array of master equations with respect to *P_m,X_*(*t*) [15, 33, 39]:

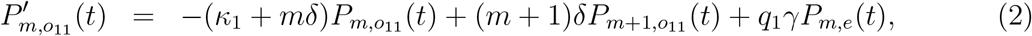

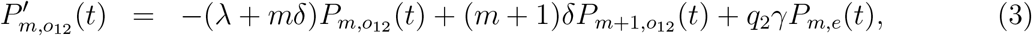

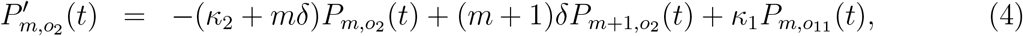

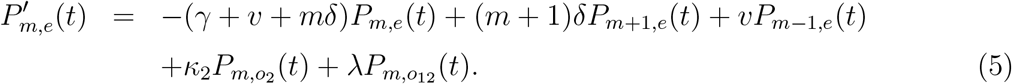

Summing up (2)-(5) gives the master equation of *P_m_*(*t*):

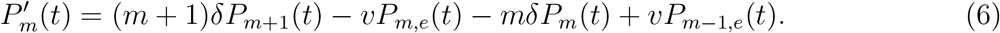

We did not focus on solving (2)-(6), which are beyond the scope of current mathematical methods within all parameter regions [22–26]. However, the master equations set a basis for calculating analytical forms for the gene state probabilities 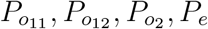, and mean transcript level *M* (*t*), defined as:

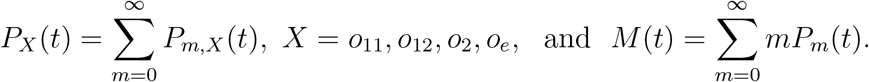

Using these definitions and (2)-(6), we derived the equations for the four state probabilities and the mean transcription level:

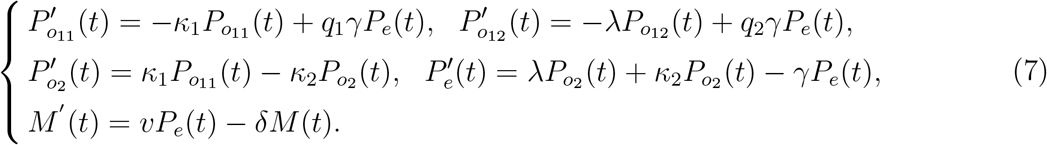

Because the system must reside on exactly one gene state at any time, we set an arbitrary initial condition for (7):

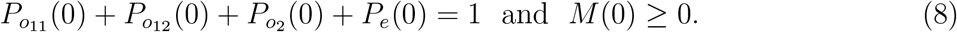

We utilized the Laplace transform method to solve the first-order differential system (7)-(8) (*Materials and Methods*). We defined a polynomial function as:

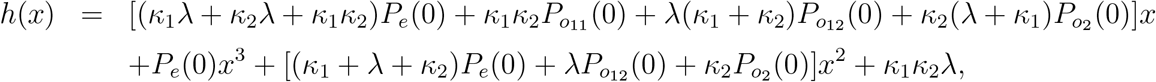

and two auxiliary numbers *c*_1_ and *c*_2_ in terms of the system parameters (*Materials and Methods* (18)-(19)). It can be verified that zero is a simple eigenvalue of the coefficient matrix for the system of the first four equations in (7). The other non-zero eigenvalues *a*_1_, *a*_2_, and *a*_3_ are calculated in terms of *c*_1_, *c*_2_, and system parameters (*Materials and Methods* (15)-(19)). Under the arbitrary initial condition (8), the average mRNA level *M* (*t*) is found to be as follows:

1. If 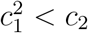, then *a*_1_, *a*_2_, *a*_3_ are real numbers with 0 < *a*_1_ < *a*_2_ < *a*_3_ (*Materials and Methods* (22)), and *M* (*t*) takes the form of:

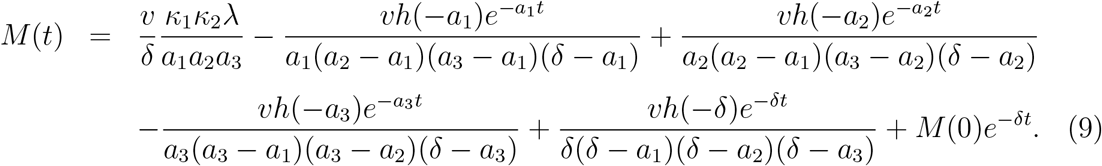
2. If 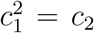, then *a*_1_, *a*_2_, *a*_3_ > 0 are real numbers with *a*_1_ /= *a*_2_ = *a*_3_ (*Materials and Methods* (24)), and *M* (*t*) takes the form of:

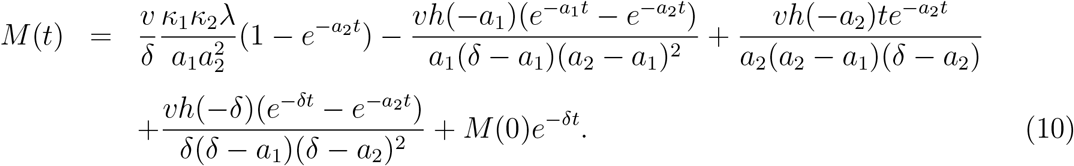
3. If 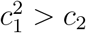, then *a*_1_ > 0 is a real number, whereas *a*_2_ and *a*_3_ are conjugate complexes, then let *a_r_* = Re(*a*_2_) and *a_i_* = Im(*a*_2_) (Materials and Methods (26)). Then, *M* (*t*) takes the form of:

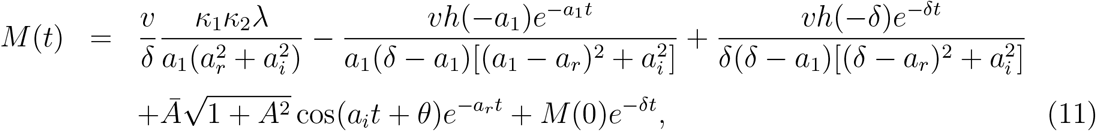

where the constants 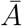, *A*, and *θ* correlate with the parameters and initial conditions (*Materials and Methods* (28)).

We characterized the dynamic profiles of *M* (*t*) with almost no expression products at the initial time *t* [11, 21, 30]. We assumed that transcription starts from the gene *off* 1 state and counts only the newly produced mRNA molecules. This gives the following initial values:

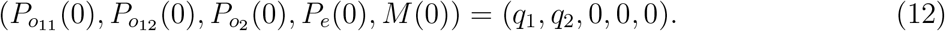

We started with an interesting case, 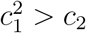. Then, *M* (*t*) was expressed in exact form (11), which contains a cosine function, and suggested possible oscillatory dynamics of *M* (*t*). However, the coefficient of the cosine function damps exponentially, which dramatically weakened the oscillation visually, as manifested by our numerical examples and observations from the three-state model [34]. The exact form (11) can only be viewed as a steady-state value, adding three exponential functions at most. If the exact form of *M* (*t*) with *M* (0) = 0 has such a structure, then it can be proved that the most complex dynamics of *M* (*t*) will only take a unique peak such as up-and-down or up-down-up profiles (*Materials and Methods* Theorem 1).

For the other case of 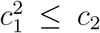, the eigenvalues *a*_1_, *a*_2_, and *a*_3_ are real, and the exact forms (9)-(10) of *M* (*t*) do not contain oscillatory functions but contain multiple exponential functions. The case 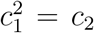 rarely occurs in biology and can be viewed mathematically as a limiting case of 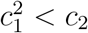. For 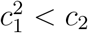, a rigorous statement of the *M* (*t*) profiles is inevitably technical. Let *x*_1_ and *x*_2_ denote the two roots of:

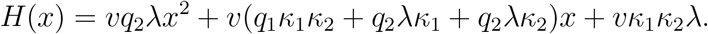

When both *x*_1_ and *x*_2_ are complex numbers, there are a total of four parameter correlations, and we showed that *M* (*t*), expressed by (9), either increases monotonically for all *t* > 0, or develops an up-down-up profile. If *x*_1_ and *x*_2_ are real values, we can classify all 60 correlations among *x*_1_, *x*_2_, *a*_1_, *a*_2_, *a*_3_, and *δ* into three categories that correspond to three distinct dynamical behaviors: the increasing, up-and-down, and up-down-up profiles of *M* (*t*). We illustrated detailed mathematical results and their proof in Theorem 2 of the *Materials and Methods*. In summary, for the cross-talking three-state model (1), even if *M* (*t*) is expressed in different exact forms (9)-(11) and influenced by various parameter correlations, *M* (*t*) can exhibit, but exhibits only three features: increasing, up-and-down, and up-down-up dynamics.

### 2.3 Fitting dynamical transcription data of mouse fibroblast genes

We demonstrated three dynamical profiles of *M* (*t*) generated by model (1). These distinct behaviors are in agreement with the observed dynamic trends of average mRNA levels in mouse fibroblast genes in response to TNF [30, 31]. For instance, Hao and Baltimore [30] divided 180 activated mouse fibroblast genes under TNF induction into three groups, separately characterized by the short, median, and long half-lives of the transcripts. As shown in Fig. 2, they found that group I genes responded quickly by forming a sharp dynamical peak at average transcription levels, group II genes did not respond quickly but most still formed a gentle transcription peak along the timeline, and group III mRNAs accumulated rather slowly and gradually increased in abundance during the observation window. The transcription data from the 12 representative mouse fibroblast genes shown in Fig. 2 contains four genes displaying the up-down-up trend of transcription dynamics (*Edn1, Cxcl1, Ccl2, Icam1*), as well as others displaying either dynamical up-and-down or monotonic increasing transcription. Using exact forms (9)-(11) of *M* (*t*), the model (1) provides good theoretical fits to all 12 datasets (coefficient of determination *R*^2^ > 0.9).

**Figure 2:**
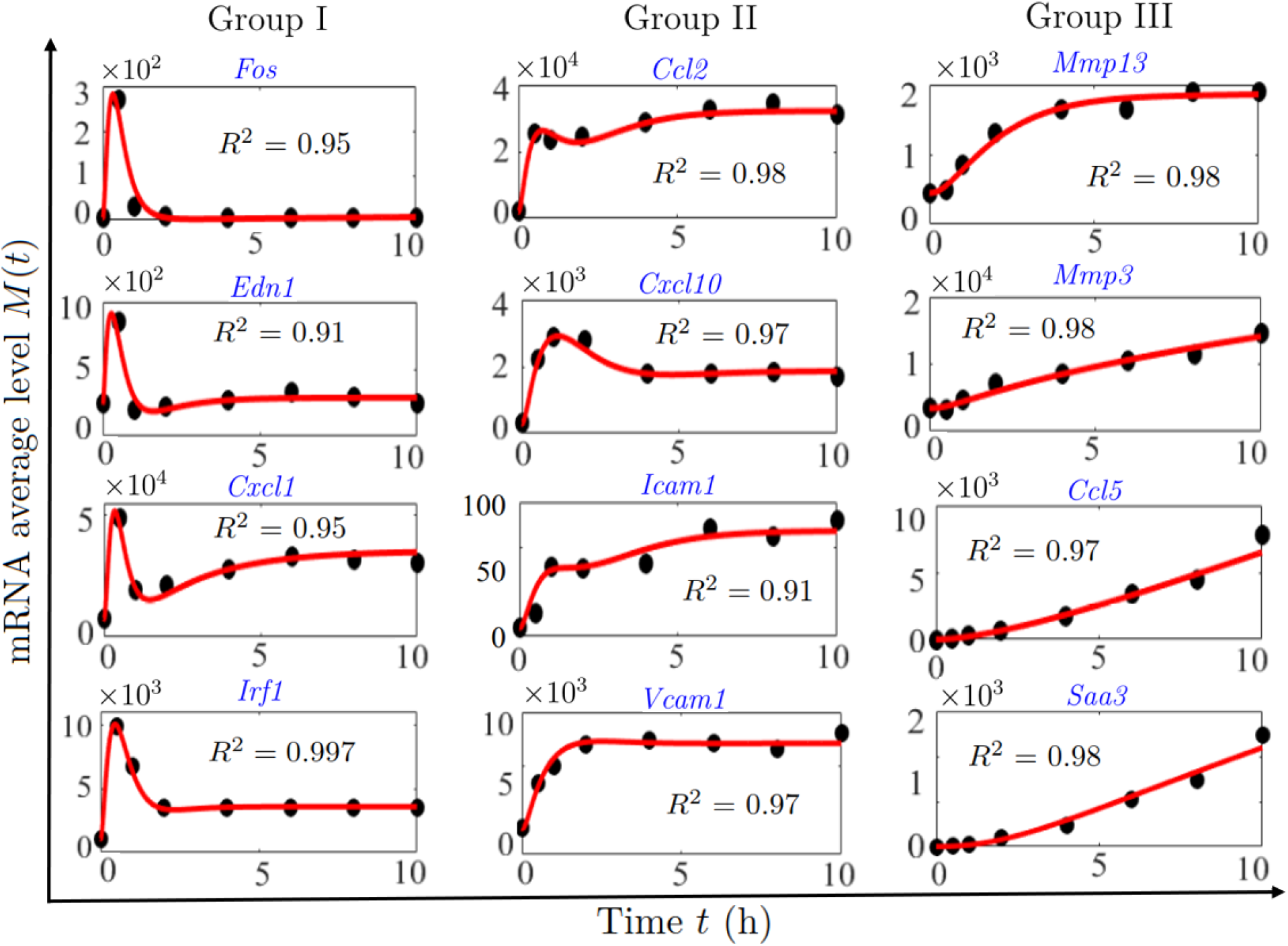
Fit of transcription data by the cross-talking three-state model (1). Black circles represent the dynamical data of mRNA average levels for 12 mouse fibroblast genes under TNF induction [30]. The genes are divided into groups I, II, and III, which are separately characterized by the short, median, and long half-lives of the transcripts [30]. The red lines are generated using exact forms (9)-(11) of the model (1), which provide a good fit to the data points (*R*^2^ > 0.9). The fitted system parameters are listed in Table S1 (*Supporting Information*).

When fitting the data shown in Fig. 2, we assigned extremely small values to the activation rates *κ*_1_ and *κ*_2_ of weak basal pathway, and restricted the mRNA degradation rate *δ* of each gene within the *δ* region of the gene group to which they belong [30] (*Supporting Information*, Table S1). Therefore, the fitted *δ* values show a significant negative correlation with gene groups I, II, and III because the gene groups themselves are classified by their transcript half-lives (Fig. 3a). Intriguingly, the freely fitted inactivation rate *γ* and activation rate *λ* of the signaling pathway also exhibited negative correlation with gene groups (Fig. 3a). This observation suggests that the simultaneous large *λ* and *γ* may separately help enhance the height of the peak and suppress the stationary values of *M* (*t*) in group I. In contrast, the small *λ* and *γ* implemented contrary functions that destroy the dynamical peak of *M* (*t*) and lift up the stationary mRNA numbers in group III. However, the probability *q*_2_ of the signaling pathway does not clearly correlate with the gene groups but varies for different genes (Fig. 3a). Note that the dynamics of *M* (*t*) in the three gene groups are mainly discriminated by its first peak. Our observations suggest that, once cells receive external cues, the frequency of the signaling pathway directing gene activation may not play a crucial role in regulating the dynamical peak of transcription level.

**Figure 3:**
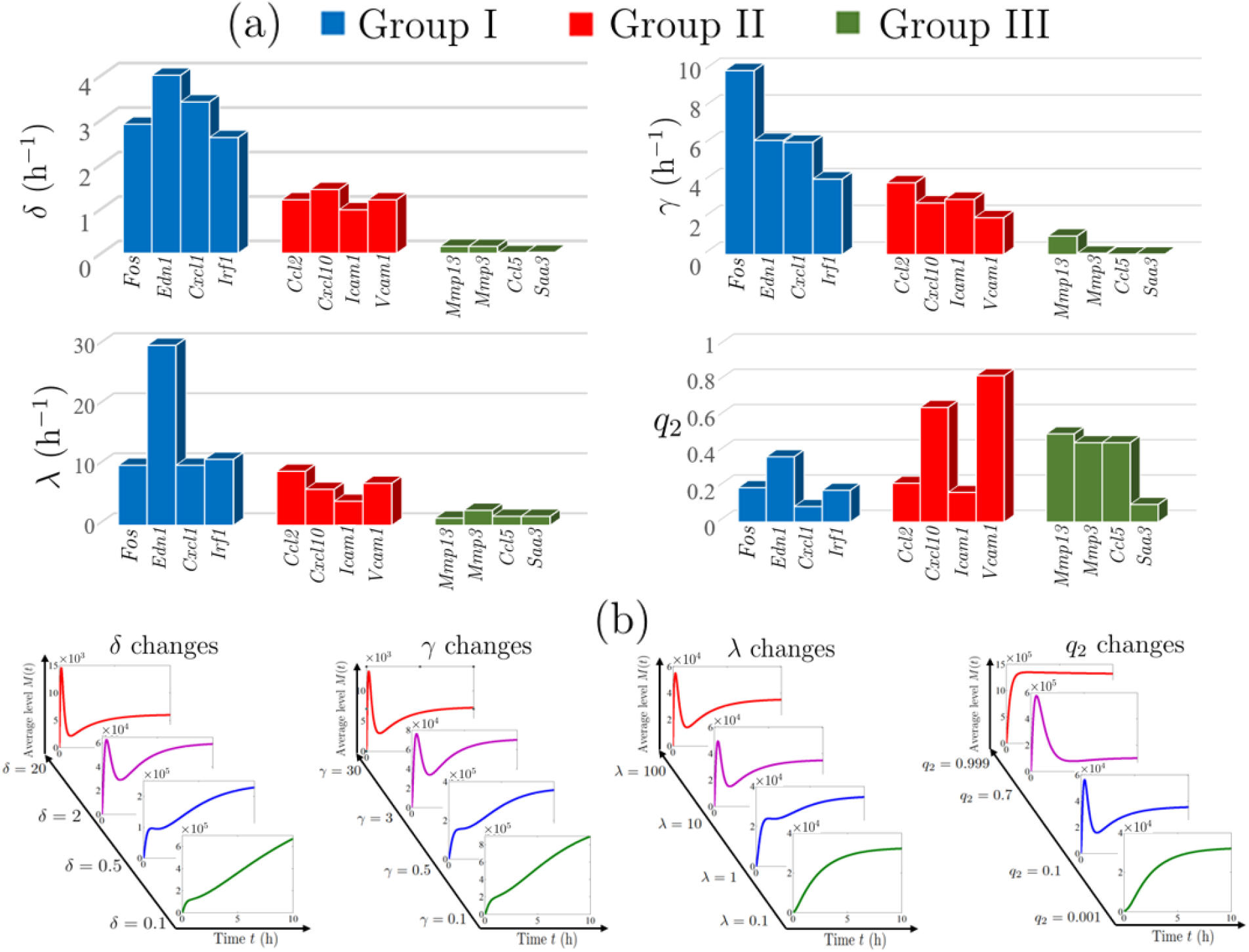
Regulation of transcription dynamics by system parameters. (a) The fitted parameters for 12 mouse fibroblast genes are listed in units of gene groups (*Supporting Information*, Table S1). Different gene groups exhibit distinct temporal transcription modes [30]. The values of mRNA degradation rate *δ*, inactivation rate *γ*, and activation rate *λ* of the signaling pathway are all negatively correlated with gene groups I, II, and III. The probability *q*_2_ of the signaling pathway does not follow a clear correlation with the gene groups. (b) Based on the fitted parameter set for the *Cxcl1* gene, increasing *δ, γ*, and *λ* transit transcription dynamics from monotonic to up-down-up mode. An increase in *q*_2_ generates switches among multiple transcription dynamical modes, where the non-monotonic modes are displayed within most of the *q*_2_ variation region (0, 1).

To further understand the regulation of *M* (*t*) profiles, we varied the parameters *δ, γ, λ*, and *q*_2_ under the fitted parameter sets of all 12 genes in Fig. 2. This procedure reveals a uniform regulation mode for each system parameter. As shown in Fig. 3b for *Cxcl1* gene, the variation in *δ* behaves as a bilateral switch to regulate *M* (*t*): there is a threshold value such that *M* (*t*) increases monotonically while *δ* stays below the threshold, but switches to a non-monotonic profile once *δ* exceeds the threshold. Such bilateral regulation of *δ* has been observed to play an important role in controlling the temporal transcription mode in mouse fibroblasts [30, 31]. In addition, both *γ* and *λ* play the same bilateral roles in regulating *M* (*t*) dynamics (Fig. 3b), reinforcing previous observation of smaller *δ, γ, λ* for group III genes that generate increasing *M* (*t*) with larger *δ, γ, λ* for groups I and II genes that exhibit non-monotonic transcription dynamics (Fig. 3a). The regulation scenario of *q*_2_ is different because it generates multiple switches among distinct *M* (*t*) profiles, when *q*_2_ increases from 0 to 1 (Fig. 3b). Exceptions need to be made for cases where *q*_2_ approaches 0 or 1, which generates increasing transcription dynamics, because the dynamical peak of *M* (*t*) seems to be robust in almost all the variation regions of *q*_2_ ∈ (0, 1). Note that *q*_2_ is closely related to signal strength. Our observations fit with the ubiquitous transcription dynamical peak of mouse fibroblast genes under TNF induction from the lowest to the highest levels [31].

### 2.4 Cross-talking *n*-state model for oscillatory transcription dynamics

Our bijection theory shows that the cross-talking three-state model (1) cannot generate multiple dynamical peaks of *M* (*t*). Therefore, the model (1) can be ruled out when *M* (*t*) exhibits oscillation [30, 32]. We focused on transcription with constant kinetic rates under stable inductions to avoid complicated *M* (*t*) dynamics regulated by time-dependent rates in response to time-varying signals [40]. The bijection theory (Table 1) shows that multiple parallel pathways induce at most one dynamical peak, and therefore introducing more parallel pathways in model (1) may not capture oscillatory *M* (*t*). We then considered decomposing the basal pathway of model (1) into multiple sequential steps, as shown in Fig. 4a for the cross-talking *n*-state model. Calculation of *M* (*t*) can follow the same procedure for model (1), which relies on solving a system of differential equations for which the coefficient matrix may have multiple pairs of eigenvalues expressed by conjugate complexes (*Materials and Methods*). According to the classical theory of ordinary differential equations, the exact forms of *M* (*t*) may contain multiple periodic cosine functions, and it is therefore plausible to visualize oscillatory dynamics.

**Figure 4:**
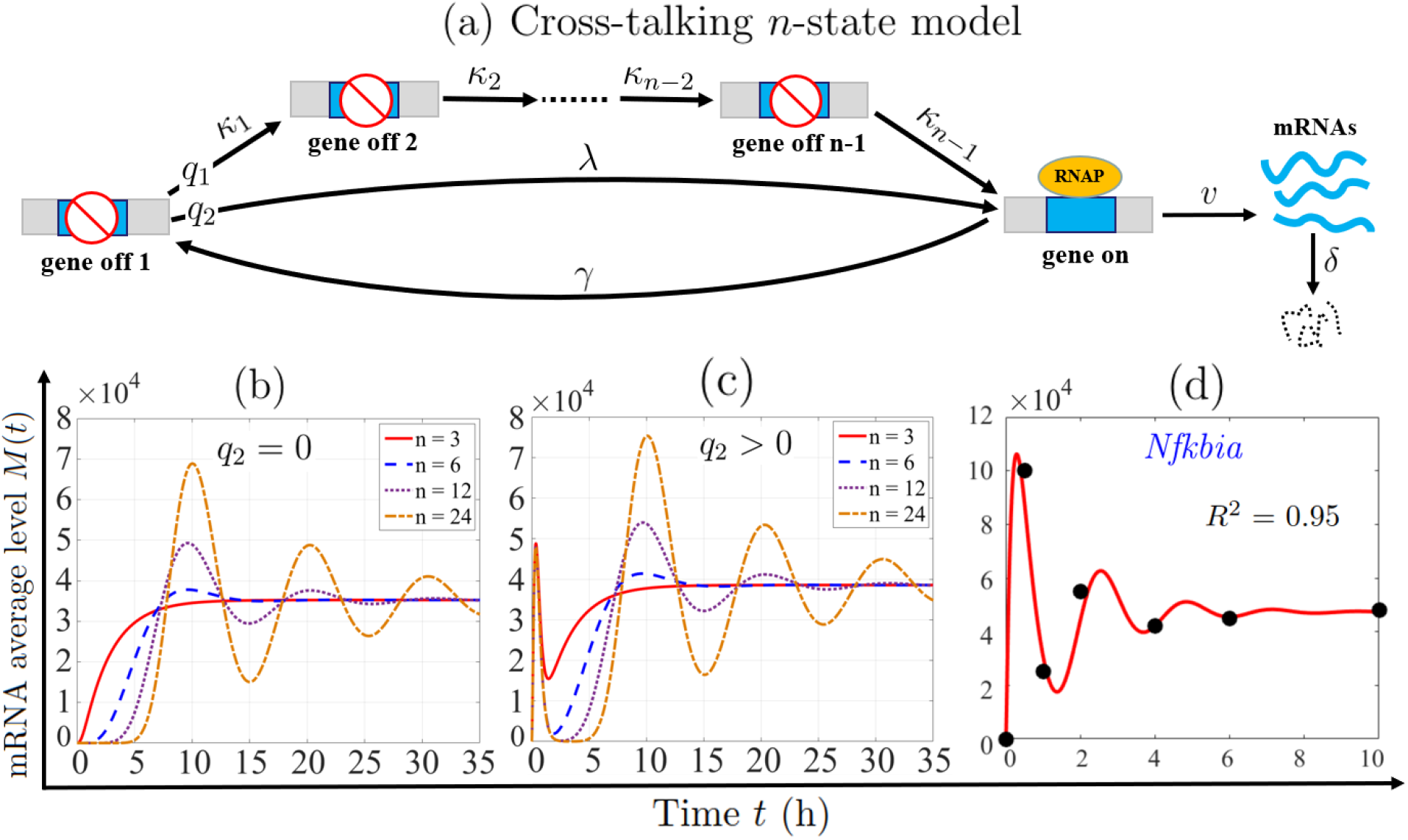
Framework for generating oscillatory transcription dynamics. (a) The cross-talking *n*-state model for which the gene is activated either by the strong signaling pathway with probability *q*_2_, or alternatively by the weak basal pathway that consists of *n* − 1 rate-limiting steps with probability *q*_1_ = 1 − *q*_2_. (b) When gene activation is directed by a single multi-step pathway (*q*_2_ = 0), the increase in the step number prolongs the initial response lag and enhances the damped oscillation of the transcription level *M* (*t*). (c) When the gene is activated by the cross-talk between pathways (*q*_2_ > 0), the increase in the step number in the basal pathway has almost no impact on the initial quick up-and-down dynamics of *M* (*t*), but significantly enhances the following damped oscillatory behavior. (d) The curve of *M* (*t*) (red lines) generated by the framework in (a) captures the oscillatory trend of transcription data (black circles) for the mouse fibroblast *Nfkbia* gene under TNF induction [30]. The parameters in (b) and (c) are the fitted rates of the *Cxcl1* gene in Table S1 with *κ*_1_ = · · · = *κ_n_*_−1_ to maintain a constant *T_off_* . Parameters in (d) are listed in Table S1.

To test *M* (*t*) oscillation induced by the sequential multi-step gene activation, we compared the *M* (*t*) profiles for different step numbers. This procedure requires that all comparisons are restricted to a constant average duration *T_off_* of the gene *off* state [15, 34]. For the cross-talk *n*-state model (Fig. 4a), *T_off_* is given by:

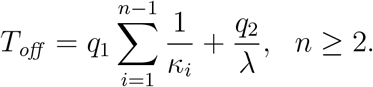

To guarantee the unchanged *T_off_* , we set three parameter scaling conditions where activation rates *κ*_1_, · · · , *κ_n_*_−1_ are separately scaled identically [14, 41], differently [8, 14], and alternatively [14]:

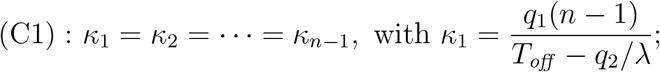

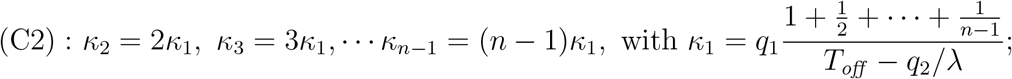

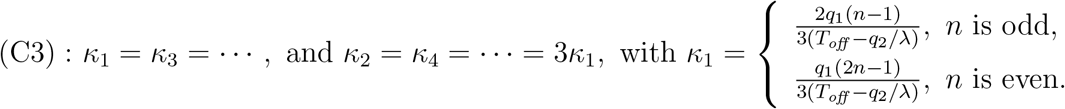

We initially examined the case of *q*_2_ = 0, for which gene activation is directed by a single multi-step pathway [14,15,26]. Under parameter scaling condition (C1), we generated several *M* (*t*) curves under different activation step numbers, using numerical simulations from the corresponding system of differential equations (*Materials and Methods* (43)). As shown in Fig. 4b under the fitted parameters of the *Cxcl1* gene in Fig. 2, multi-step gene activation for large step numbers triggers significant damped oscillatory *M* (*t*), where the significance of the oscillation is positively correlated with the step number. However, the system displays lag times of more than 8 h to reach the first peak of *M* (*t*). This slow transcription response contradicts the rapid peak of *M* (*t*) within 0.5 ∼ 2 h for mouse fibroblast genes [30, 31]. Moreover, the damped oscillation and response lag of *M* (*t*) were robust against the parameter scaling conditions (C2) and (C3) (*Supporting Information*, Fig. S1). Taken together, the large number of sequential gene activation steps facilitates the oscillatory dynamics of *M* (*t*), while a single multi-step pathway was not able to induce quickly-peaked transcription dynamics of mouse fibroblast genes.

We then quantified *q*_2_ /= 0 to introduce the cross-talking regulation of pathways on *M* (*t*) dynamics. We generated *M* (*t*) curves (*Materials and Methods* (42)) under the fitted parameters of the *Cxcl1* gene in Fig. 2 and conditions (C1)-(C3). As shown in Fig. 4c and Fig. S2 in *Supporting Information*, the system generates oscillatory *M* (*t*) with two major features. Firstly, *M* (*t*) displays a quick and sharp first peak within the initial time region, while the height and sharpness of the first peak are nearly independent of the step number and parameter scaling conditions of the basal pathway. Secondly, *M* (*t*) displays a damped and gentle second and following peaks that are tightly correlated with the step number and parameter conditions, similarly to the oscillation of *M* (*t*) induced by a single multi-step pathway (Fig. 4b). These two features capture multiple dynamical peaks for the transcription of the mouse fibroblast *Nfkbia* gene [30] (Fig. 4d). Taken together, the cross-talk between signaling and basal pathways plays a dominant role in generating the first quick up-and-down transcription dynamics, while the subsequent gentle and damped transcription oscillation is induced by the multi-step regulation in the weak basal pathway.

### 2.5 Cross-verification for monotonic transcription dynamics

Our method requires rich *M* (*t*) data dynamics to rule out frameworks that cannot sufficiently capture the dynamic features. When *M* (*t*) displays simple monotonicity, all frameworks (Fig. 1) may provide good fits to the *M* (*t*) data. However, we cannot claim that the two-state model is the best model to explain the data, even if it is the simplest compared to the other frameworks. This is because monotonic *M* (*t*) may be generated by the manipulated small degradation rate *δ* (Fig. 3b) for accurately counting transcript numbers in single-cell measurements [11,14]. One way to avoid the masking effect of small *δ* on *M* (*t*)-rich dynamics is to consider the average level of nascent RNA that is independent of mRNA degradation [4,42]. In contrast, some single-cell data for transcription of mRNA molecules display smooth trend lines along the timeline, such as the noise *CV*^2^(*t*), Fano factor *ϕ*(*t*), and probability *P*_0_(*t*) of the gene producing zero transcripts. These indexes may be used in conjunction with *M* (*t*) to cross-verify the gene activation frameworks.

The calculation of *ϕ*(*t*) and *CV* ^2^(*t*) are standard, and require solving the second moment *μ*_2_(*t*) of the system of ordinary differential equations derived from the corresponding master equations [33, 39]. We calculated the exact forms of *μ*_2_(*t*) for the cross-talking three-state model (1) (*Materials and Methods* (48)), and *ϕ*(*t*) = *μ*_2_(*t*)*/M* (*t*) − *M* (*t*) and *CV* ^2^(*t*) = *ϕ*(*t*)*/M* (*t*) were then readily obtained. The probability *P*_0_(*t*) is one of the solutions of master equations. Calculation of *P*_0_(*t*) involves introducing the generating function *V* (*z, t*) to transform the master equation into a system of partial differential equations for *V* (*z, t*). However, while solving *V* (*z, t*) is somewhat difficult, we can still express *V* (*z, t*) in closed forms of hypergeometric functions for different models [23, 26]. Consequently, *P*_0_(*t*) can be obtained by *P*_0_(*t*) = *V* (−1, *t*). The expressions of *ϕ*(*t*), *CV* ^2^(*t*), and *P*_0_(*t*) are not as neat as that of *M* (*t*), which prevents us from establishing theoretical bijections between their dynamical profiles and parameter regions. Nevertheless, the computation of *ϕ*(*t*), *CV* ^2^(*t*), and *P*_0_(*t*) through their expressions or the corresponding differential equations (*Materials and Methods*) can generate dynamical curves for fitting data [11, 40].

We hypothesized that different gene activation frameworks can generate distinct dynamics of *ϕ*(*t*), *CV* ^2^(*t*), and *P*_0_(*t*) under the same monotonic *M* (*t*). To verify this, we first fitted a group of increasing *M* (*t*) data to all four frameworks in Fig. 1 (see Fig. 5a). Under the fitted parameters we separately generated dynamical curves of *ϕ*(*t*), *CV* ^2^(*t*), and *P*_0_(*t*) corresponding to each framework. The curves of noise *CV* ^2^(*t*) clustered within a low-value region, probably because of their large denominator *M* (*t*) (Fig. 5b), suggesting that the noise data may not help discriminate gene activation frameworks. The curves of the Fano factor *ϕ*(*t*) are clearly separated into two categories: the curves are very high under cross-talking pathways, but are relatively low under a single pathway (Fig. 5c). This separation reinforces the conclusion that the parallel pathways enhance *ϕ* [39], while sequential steps suppress *ϕ* [15] under the same mean level *M* at steady-state. The curves of *P*_0_(*t*) follow the same separation as that of *ϕ*(*t*) (Fig. 5d), which reinforces the observed suppressed *P*_0_(*t*) under multi-step gene activation [26]. Taken together, the dynamics of *ϕ*(*t*) and *P*_0_(*t*) may help discriminate the activation frameworks regulated by cross-talking pathways or a single pathway.

**Figure 5:**
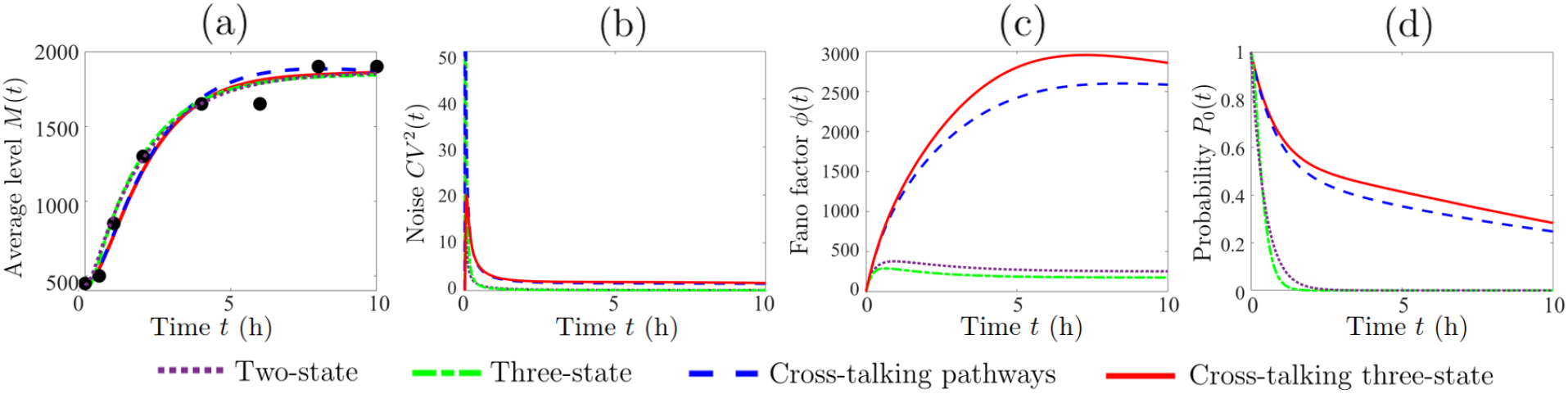
Discrimination of gene activation frameworks under monotonic dynamical mRNA average level. (a) The increasing transcription mean data (black circles, *Mmp13* gene [30]) are fitted by different models. (b) Dynamical curves of transcription noise for each model are clustered with relatively small values most of the time. For (c) transcription Fano factor and (d) probability of genes not producing any transcript, the dynamical curves generated by a single pathway deviate from the curves generated by cross-talking pathways. Parameters: *v* = 2630 h^−1^, *q*_2_ = 0.5; two-state model, (*λ, γ, δ*) = (2.21, 4, 0.5) h^−1^; three-state model, (*κ*_1_, *κ*_2_, *γ, δ*) = (5.09, 5.09, 4, 0.55) h^−1^; cross-talking pathways model, (*κ, λ, γ, δ*) = (0.07, 1.2, 1.2, 0.15) h^−1^; cross-talking three-state model, (*κ*_1_, *κ*_2_, *λ, γ, δ*) = (0.24, 0.1, 1.2, 1, 0.15) h^−1^.

## 3 Conclusion and Discussion

The classical two-state model used in single-cell studies posits that a gene will randomly transition between *on* (active) and *off* (inactive) states, with mRNA molecules being produced only when the gene is *on* (Fig. 1a). Compared to the universal feature of a single rate-limiting step turning gene *off* [6, 8, 13], the process of turning a gene *on* is usually non-Markovian, and is influenced by multiple rate-limiting fluctuations [6,7,13]. The frame-work of gene activation has been modeled by listing rate-limiting steps sequentially [8, 14] (Fig. 1b), parallelly [16, 35] (Fig. 1c), or in the form of their combinations [17] (Fig. 1d). Recent studies have facilitated efficient computation of downstream transcription distribution under non-Markovian gene activation [18, 19]. However, the best method for mapping the transcription distribution data back to the exact picture of the activation framework for the gene of interest remains elusive.

Distinct gene activation frameworks cannot be discriminated by the steady-state single-cell data on mRNA distribution or its noise, Fano factor, and mean, because the limited steady-state diagrams can be reliably illustrated by different frameworks [15, 23, 27–29]. When the time dimension is included, the distribution histograms of both gene *off* periods and mRNA copy numbers provide a complete static basis to computationally search the optimal number of rate-limiting steps in gene activation and their kinetic parameters [9, 14, 20]. However, these methods require a prior hypothetical framework of gene activation that cannot be easily deduced from limited gene *off* distribution modes [15–17] or from similar dynamical transition patterns of mRNA distribution profiles [24–26]. Therefore, an efficient method for determining confident gene activation frameworks is urgently required prior to any attempt to fit single-cell dynamical transcription data.

In this paper, we demonstrated that the dynamics of transcript average level *M* (*t*) can serve as a competent candidate to facilitate the efficient estimation of the gene activation framework and system parameters. Firstly, compared with time-consuming single-cell measurements [11,21], the dynamic features of *M* (*t*) for a large number of genes can be captured relatively quickly by traditional cell population methods. Secondly, compared to the multiple uneven distribution profiles at discrete time points [9, 20], a single smooth curve of *M* (*t*) along the timeline presents easily discriminated dynamic features and provides a more efficient theoretical fit to the data. Thirdly, compared to the lack of a method to calculate dynamical mRNA distribution [22, 23], the exact forms of *M* (*t*) are rather neat and can be achieved by the standard theory of ordinary differential equations.

By taking advantage of the *M* (*t*) expressions and their simple forms (Eqs. (9)-(11)), we were able to establish theoretical bijections between the *M* (*t*) dynamics and parameter regions for different gene activation frameworks. Subsequently , frameworks that cannot capture the exhibited *M* (*t*) dynamical features can be ruled out, while the optimal forms of the other potential frameworks are further determined by fitting *M* (*t*) data. We illustrated this idea by analyzing activation frameworks for a large number of TNF-induced mouse fibroblast genes. These genes display rich transcription dynamics that can be categorized into three main features: increasing, up-and-down, and up-down-up profiles of *M* (*t*) [30,31]. Our bijection theories (Table 1 and *Materials and Methods* Theorems 1–2) show that these three distinct dynamics cannot be achieved by the sequential or parallel rate-limiting steps alone, but can be captured by the simplest form of combined sequential and parallel steps (Fig. 1d). We call this framework the cross-talking three-state model, as depicted in (1), for which the gene is activated either by the weak basal pathway consisting of two sequential steps, or alternatively by the strong signaling pathway.

The cross-talking three-state model (1) can serve as a reliable model because it provides a good theoretical fit for all the representative datasets on the transcription dynamics of mouse fibroblast genes [30] (Fig. 2). Moreover, analysis of the freely fitted system parameters (Table S1) reveals two regulation scenarios for *M* (*t*) dynamics (Fig. 3). For the mRNA degradation rate *δ*, inactivation rate *γ*, and activation rate *λ* of the signaling pathway, *M* (*t*) maintains a monotonic increase when *δ, γ*, and *λ* remain below their thresholds while switching non-monotonically once *δ, γ*, and *λ* exceed the threshold values (Fig. 3b). This bilateral regulation results in an interesting phenomenon: *δ, γ*, and *λ* are simultaneously large for the genes displaying non-monotonic *M* (*t*), whereas they are simultaneously small for genes displaying monotonic *M* (*t*) (Fig. 3a). In contrast to the gene-specific *δ, γ*, and *λ*, the environment influenced the probability *q*_2_ that the signaling pathway does not play a crucial role in regulating the first peak of *M* (*t*) (Fig. 3). This observation is consistent with the robust transcription dynamical peak of mouse fibroblast genes, regardless of how the TNF induction level varies [31].

We note that a small number of mouse fibroblast genes display damped transcription oscillation, with the first peak forming rapidly within the initial period [30, 31]. We suggest that multiple sequential rate-limiting steps may play a role in triggering oscillatory behavior. Therefore, we developed the cross-talking *n*-state model by decomposing the basal pathway of model (1) into multiple steps (Fig. 4a). We first ruled out the framework of a single multi-step pathway as it triggers a long lag reaching the first peak of *M* (*t*) (Fig. 4b, Fig. S1), which contradicts the observed rapid peak of *M* (*t*). However, when we recovered the cross-talk between the signaling pathway and the multi-step basal pathway, the initial lag time disappeared and the first peak of *M* (*t*) formed quickly (Fig. 4c, Fig. S2). Intriguingly, the step number and parameter scaling condition in the basal pathway have almost no effect on the first peak of *M* (*t*), whereas they significantly influence the other dynamical peaks. Together with the transcription data of the mouse fibroblast genes (Fig. 4d), we found that cross-talking regulation between pathways is crucial to trigger the first rapid, sharp peak of *M* (*t*), while the multi-step regulation facilitates the following damped and gentle oscillatory dynamics.

Thus, we developed a procedure to estimate frameworks of gene activation using transcription level *M* (*t*) dynamical data at the cell population level. This procedure readily disqualifies frameworks that do no perform satisfactorily, and determines potential frameworks that provide the best data fit. When more sophisticated single-cell distribution data of the gene *off* period [8,14] or mRNA copy numbers [9,20,21] are used, our procedure provides a prior estimation of activation frameworks and parameter rates to facilitate more computationally efficient search for optimal step numbers and parameters. This procedure relies on the rich dynamics of *M* (*t*). When *M* (*t*) behaves monotonically, the other transcription indexes are required to cross-verify the activation frameworks. For instance, the dynamical Fano factor or the probability of no transcript being produced may help discriminate frameworks regulated by a single pathway or cross-talking pathways (Fig. 5).

Future work may utilize additional data for different genes and transcription indexes to test and develop our procedure. In addition, we anticipate the inclusion of nascent RNA data that are not influenced by the mRNA degradation rate and may therefore provide more direct information on gene activation frameworks [4,42]. Finally, a cell cycle description [22,41,43] may be introduced into gene activation frameworks to eliminate estimation errors caused by disregarding cell cycle stochasticity [41].

## Materials and Methods

### Exact forms of *M* (*t*) for cross-talking three-state model (1)

To calculate mRNA average level *M* (*t*) for the model (1), we need solve the first four differential equations in system (7) for gene state probabilities 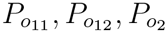 and *P_e_* under the arbitrary initial condition (8):

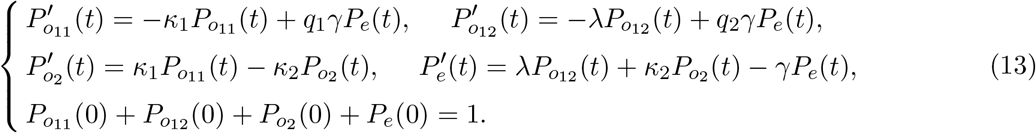

To solve the above first-order differential system, we denote by

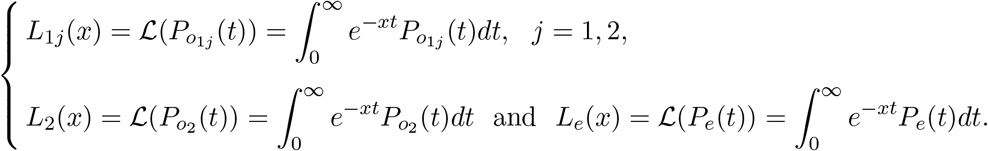

Then using Laplace transform in the theory of ordinary differential equation to system (13) yields

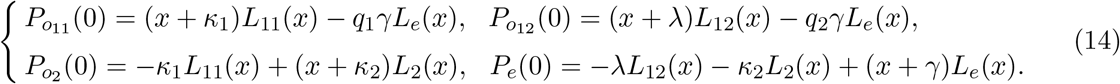

The determinant of coefficient matrix for the above system can be directly calculated as

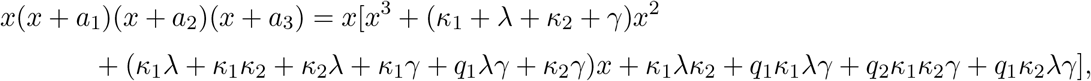

where *a*_1_, *a*_2_ and *a*_3_ are non-zero eigenvalues of the coefficient matrix for the system (13). To calculate *a*_1_, *a*_2_ and *a*_3_, we let the above formula equal to 0, and then solve the cubic polynomial equation using Cardano formula. This gives

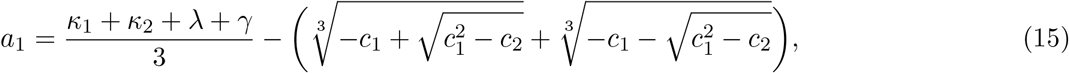

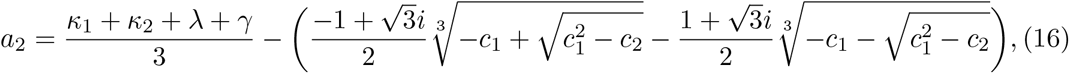

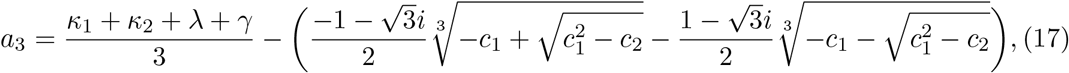

where

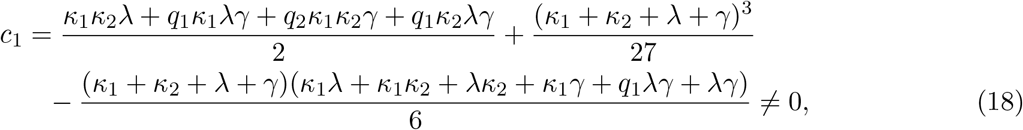

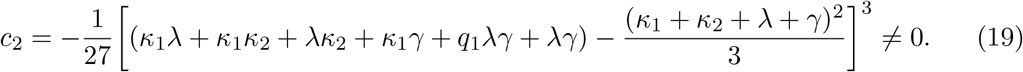

Applying Cramer’s rule in the theory of linear algebra on the system (14) readily gives

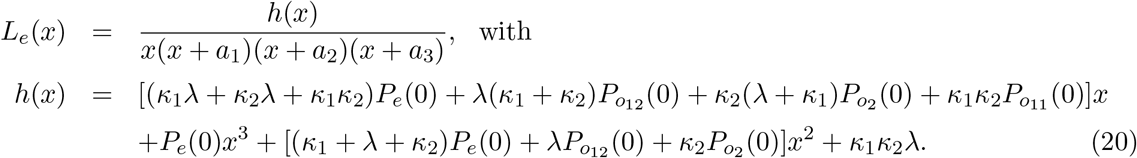

To calculate analytical formula for mRNA average level *M* (*t*), we first notice that it satisfies

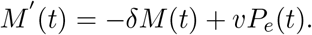

Then applying Laplace transform to *M* (*t*) gives

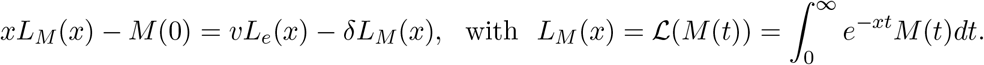

The substitution of *L_e_*(*x*) in (20) further leads to

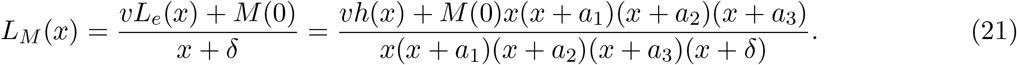

1. If 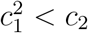, then *a*_1_, *a*_2_ and *a*_3_ given in (15)-(19) are real numbers satisfying 0 < *a*_1_ < *a*_2_ < *a*_3_. Also, the direct calculation can simplify *a*_1_, *a*_2_ and *a*_3_ to be

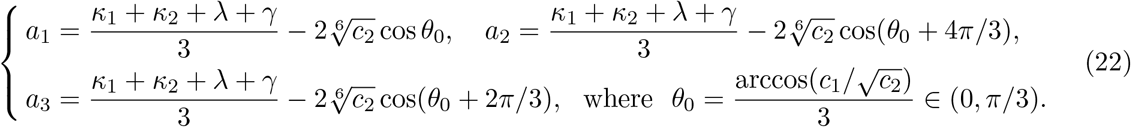 We decompose *L_M_* (*x*) given by (21) into the sum of partial fractions as

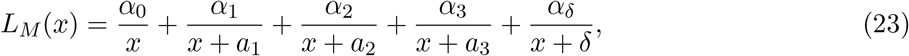

where the coefficients *α*_0_, *α*_1_, *α*_2_, *α*_3_ and *α_δ_* satisfy

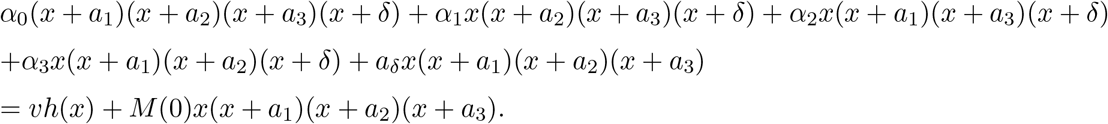 The separate substitution of *x* = 0, −*a*_1_, , −*a*_2_, −*a*_3_, −*δ* into above equation yields

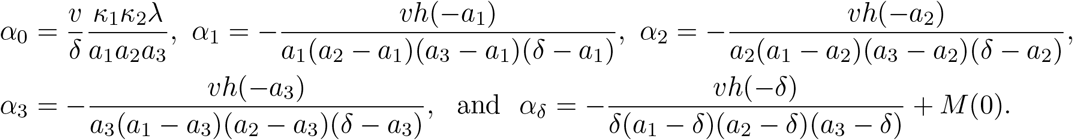 Then the application of the inverse Laplace transform to *L_M_* (*x*) given in (23) readily leads to

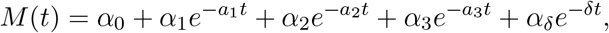

which is exactly the expression (9).
2. If 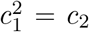, then it can be verified that *a*_1_, *a*_2_ and *a*_3_ given by (15)-(19) are positive real numbers taking the form of

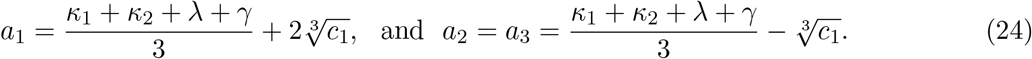 We decompose *L_M_* (*x*) given in (21) into the sum of partial fractions as

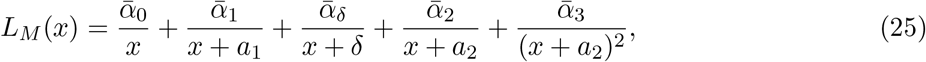

where the coefficients 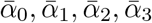 and 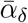 satisfy

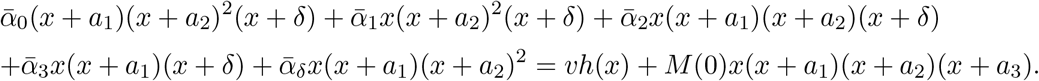 Then the separate substitution of *x* = 0, −*a*_1_, −*a*_2_, −*δ* into this equation yields

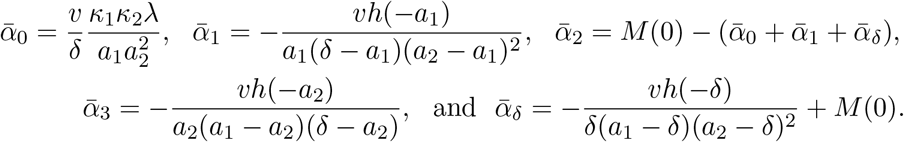 Then the application of the inverse Laplace transform to *L_M_* (*x*) given in (25) readily leads to

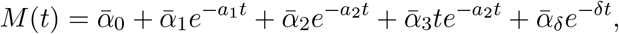

which verifies the expression (10).
3. If 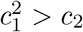, then *a*_1_ is a positive real number while *a*_2_ and *a*_3_ are conjugate complexes. Let *a_r_* = Re(*a*_2_) and *a_i_* = Im(*a*_2_). Then expressions (15)-(19) suggest

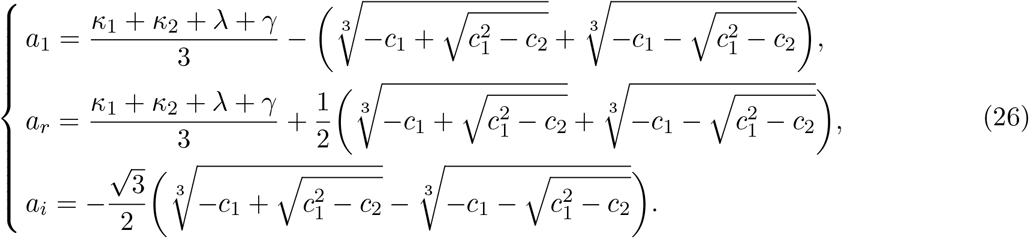 We decompose *L_M_* (*x*) of (21) into the sum of partial fractions as

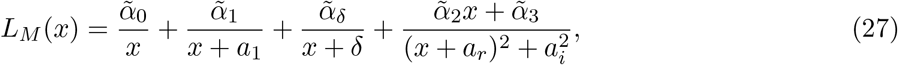

with 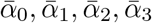 and 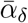 satisfying

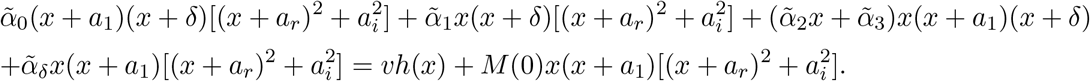 Then the separate substitution of *x* = 0, −*a*_1_, −*δ* into this equation yields

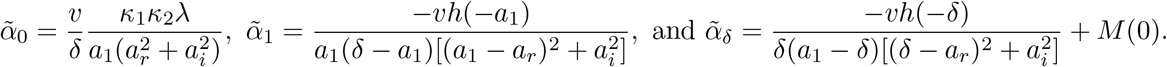 The other two 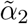 and 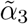 are obtained by comparing the coefficients:

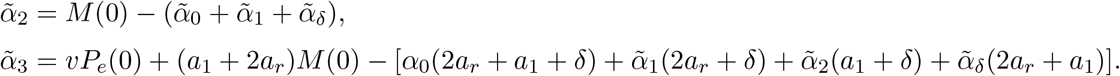 Using above 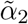 and 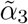 we further introduce the symbols

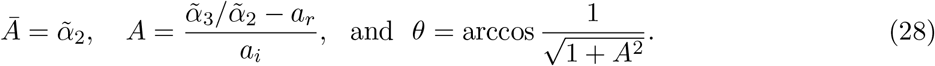 Then the application of inverse Laplace transform to *L_M_* (*x*) given in (27) readily leads to

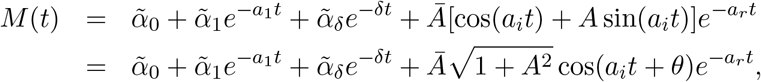

which is exactly the expression (11).

### Dynamical profiles of *M* (*t*) for cross-talking three-state model (1)

In this section we utilize exact forms of *M* (*t*) to understand its dynamical features. We consider the initial condition (12) for which the gene is silence and there is almost no transcripts at the initial time [11, 21, 30]. The following two theorems illustrate the cases of 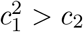 and 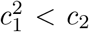, respectively.

#### Theorem 1.

For arbitrary non-zero real numbers a, b, c and positive real numbers *M* ^∗^, α, β, r, if M (t) is expressed as

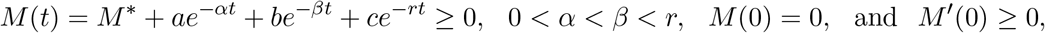

then M (t) displays only one of increasing, up-and-down and up-down-up dynamical profiles.

**Proof.** The derivative of *M* (*t*) gives

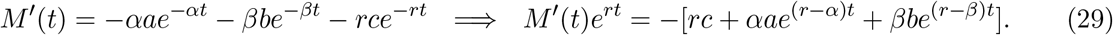

We divided the discussion into two cases of *a* > 0 and *a* < 0.

I. If *a* > 0, then lim_*t*→∞_ *M*′^t^(*t*) < 0 by noting that the sign of *M* ^t^(*t*) is dominated by the term “ − *αae*^−*αt*^” when *t* → ∞. Taking the derivative of *M* ^t^(*t*)*e^rt^* in (29) gives

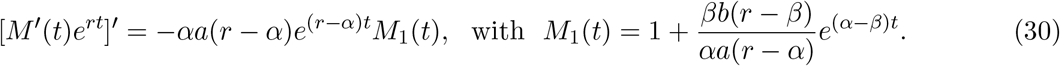 If *b* ≥ −*αa*(*r* − *α*)*/β*(*r* − *β*), then (30) indicates

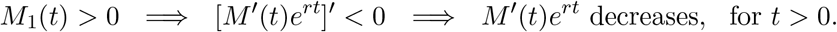 There are two probabilities of *M*′^t^(0) > 0 and *M*′^t^(0) = 0. If *M*′^t^(0) > 0, then together with lim_*t*→∞_ *M*′^t^(*t*) < 0, there exists *t*_1_ > 0 such that

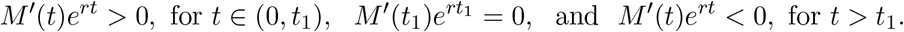 Therefore, *M* (*t*) increases for *t* ∈ (0, *t*_1_) and decreases for *t > t*_1_ which presents up-and-down dynamics. If *M* ^t^(0) = 0, then *M* ^t^(*t*)*e^rt^* < 0 for *t* > 0, and thus *M* (*t*) decreases with *M* (*t*) < 0 all the time. This contradicts to the assumption of *M* (*t*) ≥ 0. If *b* < −*αa*(*r* − *α*)*/β*(*r* − *β*), then (30) suggests that there is *t*_2_ > 0 such that

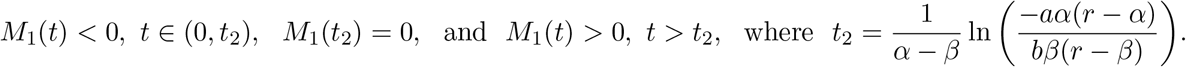 Then (30) suggests that [*M*′^t^(*t*)*e^rt^*]′ > 0 for *t* ∈ (0, *t*_2_) and [*M*′(*t*)*e^rt^*]′ < 0 for *t > t*_2_. Therefore, *M*′(*t*)*e^rt^* increases in (0, *t*_2_) while decreases in (*t*_2_, ∞). Since *M*′(0) ≥ 0 and lim_*t*→∞_ *M*′(*t*) < 0, there exists *t*_3_ such that 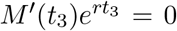. Thus *M* ^t^(*t*)*e^rt^* > 0 for *t* ∈ (0, *t*_3_) and *M* ^t^(*t*)*e^rt^* < 0 for *t > t*_3_. This indicates that *M* (*t*) increases for *t* ∈ (0, *t*_3_) and decreases for *t > t*_3_, which is the up-and-down dynamics.
II. If *a* < 0, the (29) indicates that lim_*t*→∞_ *M*′(*t*) > 0. If *b* ≤ −*αa*(*r* − *α*)/*β*(*r* − *β*), then (30) gives

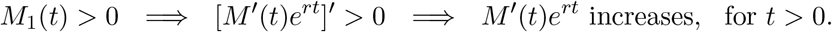

Since *M*′ (0) ≥ 0, we have *M*′ (*t*)*e^rt^* ≥ 0, *t* > 0 which suggests that *M* (*t*) increases for all the time.

On the other hand, if *b* > −*αa*(*r* − *α*)/*β*(*r* − *β*), then from (30) we find that there exists *t*_4_ > 0 such that

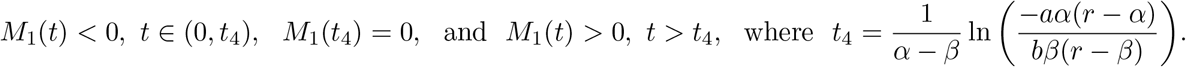

Together with (30), this further leads to [*M*′ (*t*)*e^rt^*]′ < 0 for *t* ∈ (0, *t*_4_) and [*M*′ (*t*)*e^rt^*]^t^ > 0 for *t* > *t*_4_. Therefore, *M*′ (*t*)*e^rt^* decreases in (0, *t*_4_) while increases in (*t*_4_, ∞). Since *M*′ (0) ≥ 0, there are two possibilities of 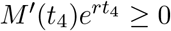 and 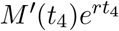 < 0. For 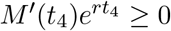, the fact of lim_*t*→∞_ *M*′ (*t*) > 0 indicates *M*′ (*t*)*e^rt^* ≥ 0 for all *t* ≥ 0, and thereby *M* (*t*) increases all the time. If it is 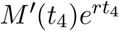 < 0, then there exist 0 < *t*_5_ < *t*_4_ < *t*_6_ such that 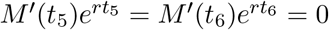 and

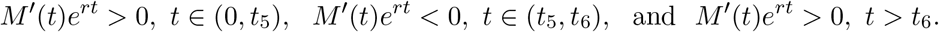

This indicates that *M* (*t*) displays an up-down-up dynamics that *M* (*t*) increases for *t* ∈ (0, *t*_5_) ∪ (*t*_6_, ∞) and decreases within *t* ∈ (*t*_5_, *t*_6_). The proof is completed.

#### Theorem 2.

Let Condition 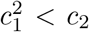 hold. Let

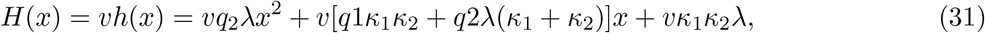

and x_1_ and x_2_ be two roots of H(x). Without loss of generality, assume δ ≠ a_1_, a_2_, a_3_ given by (22). If x_1_ and x_2_ are complex valued, then either M (t) increases monotonically for all t > 0, or M (t) develops up-down-up dynamics. If x_1_ and x_2_ are real valued and x_1_ < x_2_, then we have:

1. If one of the following occurs: (i) −x_2_ < {a_1_, a_2_, a_3_, δ} < −x_1_, (ii) −x_2_ < {a_1_, a_2_, δ} < −x_1_ < a_3_, (iii) −x_2_ < {a_1_, δ} < −x_1_ < a_2_, (iv) −x_2_ < a_1_ < a_2_ < a_3_ < −x_1_ < δ, (v) −x_2_ < a_1_ < a_2_ < −x_1_ < min{a_3_, δ}, (vi) −x_2_ < a_1_ < −x_1_ < min{a_2_, δ}, (vii) −x_2_ < δ < −x_1_ < a_1_, then m(t) increases initially until reaching a peak and then goes down (up-and-down).
2. If one of the following occurs: (i) a_1_ < −x_2_ < a_2_ < a_3_ < −x_1_ < δ, (ii) a_1_ < −x_2_ < a_2_ < a_3_ < −x_1_ < δ, (iii) a_1_ < −x_2_ < a_2_ < −x_1_ < min{a_3_, δ}, (iv) a_1_ < −x_2_ < {a_2_, δ} < −x_1_ < a_3_, (v) max{a_1_, δ} < −x_2_ < a_2_ < −x_1_ < a_3_, (vi) δ < −x_2_ < a_1_ < a_2_ < −x_1_ < a_3_, (vii) a_1_ < −x_2_ < δ < −x_1_ < a_2_, and (viii) δ < −x_2_ < a_1_ < −x_1_ < a_2_, then M (t) increases monotonically for all t > 0.
3. If one of the following occurs: (i) −x_1_ < min{a_1_, δ}, (ii) a_1_ < −x_2_ < −x_1_ < min{a_2_, δ}, (iii) a_2_ < −x_2_ < −x_1_ < min{a_3_, δ}, (iv) a_3_ < −x_2_ < −x_1_ < δ, (v) max{a_1_, δ} < −x_2_, (vi) max{a_2_, δ} < −x_2_ < −x_1_ < a_3_, (vii) max{a_3_, δ} < −x_2_, (viii) δ < −x_2_ < −x_1_ < a_1_, (ix) a_3_ < −x_2_ < δ < −x_1_, (x) a_2_ < −x_2_ < a_3_ < −x_1_ < δ,(xi) a_2_ < −x_2_ < {a_3_, δ} < −x_1_, (xii) {a_2_, δ} < −x_2_ < a_3_ < −x_1_, (xiii) a_1_ < −x_2_ < {a_2_, a_3_, δ} < −x_1_, (xiv) {a_1_, δ} < −x_2_ < a_2_ < a_3_ < −x_1_, (xv) δ < −x_2_ < a_1_ < a_2_ < a_3_ < −x_1_, then either M (t) increases monotonically for all t > 0, or M (t) develops up-down-up dynamical profile.

**Proof.** The substitution of the initial condition (12) into the differential system (7) gives *M*′ (0) = 0, 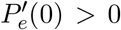, and in turn, 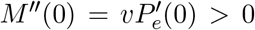. Consequently, *M* (*t*), *M*′ (*t*), and *M*” (*t*) are all positive for *t* > 0 sufficiently small. The initial values (12) simplifies the exact form (9) of *M* (*t*) in the form of

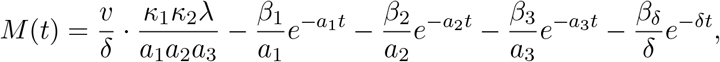

where

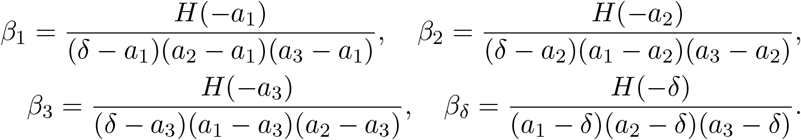

Thus

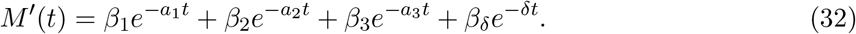

We first consider the situation of complex valued *x*_1_ and *x*_2_. There are total 4 parameter cases and we only present the proof for the case *δ* < *a*_1_ < *a*_2_ < *a*_3_, since the other cases *a*_1_ < *δ* < *a*_2_ < *a*_3_, *a*_1_ < *a*_2_ < *δ < a*_3_ and *a*_1_ < *a*_2_ < *a*_3_ < *δ* can be treated similarly. If *x*_1_ and *x*_2_ are complex valued, then *H*(*x*) > 0 for all *x*. In view of *δ* < *a*_1_ < *a*_2_ < *a*_3_, we obtain

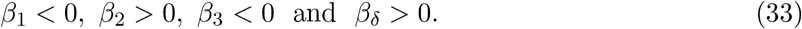

For *t* > 0 sufficiently large, *M*′ (*t*) is dominated by *β_δ_* exp(−*δt*) in (32) and is therefore positive. It follows from (32) that

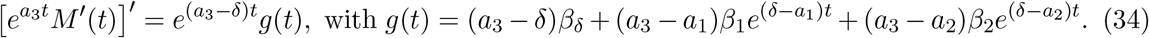

It is seen that

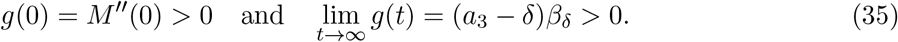

We can see from (34) that

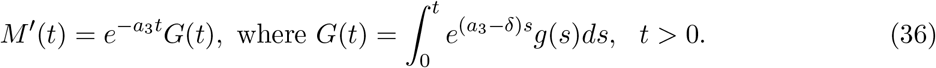

It follows from (32) and (36) that

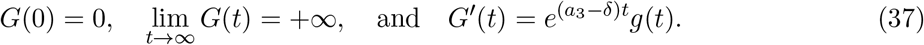

Differentiating *g*(*t*) gives

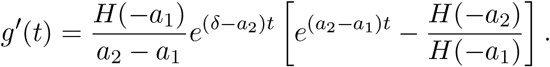

If *H*(−*a*_2_) ≤ *H*(−*a*_1_), then *g*′(*t*) ≥ 0, and so *g*(*t*) > 0 for all *t* > 0. Hence (36) gives *M*′ (*t*) > 0 for all *t* > 0. If *H*(−*a*_2_) > *H*(−*a*_1_), then *g*′(*t*) < 0 in (0, *t*_0_) for

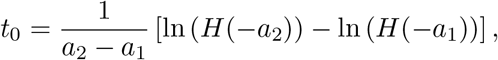

and becomes positive for *t* > *t*_0_. If *g*(*t*_0_) ≥ 0, then *g*(*t*) > 0 for all *t* > 0 and *t* ≠ *t*_0_. Thus (36) gives again *M*′ (*t*) > 0 for all *t* > 0. If *g*(*t*_0_) < 0, then (35) implies that *g*(*t*) has two zeros *t*_1_ > 0 and *t*_2_ > 0 with *g*(*t*) > 0 in (0, *t*_1_) ∪ (*t*_2_, +∞), and *g*(*t*) < 0 in (*t*_1_, *t*_2_). It follows from (37) that

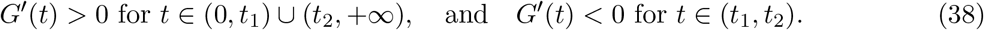

If *G*(*t*_2_) ≥ 0, then *G*(*t*) > 0 for all *t* > 0 and *t* ≠ *t*_2_, and so *M*′ (*t*) > 0 for all *t* > 0 and *t* ≠ *t*_2_. We recall that *G*(*t*) > 0 for *t* > 0 both sufficiently small and large. Thus, if *G*(*t*_2_) < 0, then (38) implies that *G*(*t*) has two zeros *t*_3_ > 0 and *t*_4_ > 0 with *G*(*t*) > 0 in (0, *t*_3_) ∪ (*t*_4_, +∞), and *G*(*t*) < 0 in (*t*_3_, *t*_4_). It follows from (36) that *M*′ (*t*) > 0 in (0, *t*_3_) ∪ (*t*_4_, +∞) and *M*′ (*t*) < 0 in (*t*_3_, *t*_4_). This finishes the proof of the case when *x*_1_ and *x*_2_ are complex valued.

In the following, we assume that *x*_1_ and *x*_2_ are real valued.

1. We present the proof with the additional assumption that −*x*_2_ < *a*_1_ < *a*_2_ < *δ* < −*x*_1_ < *a*_3_, as the other cases can be dealt with by the same idea. Since

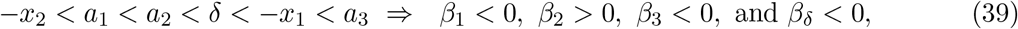

by using (32) again, we obtain

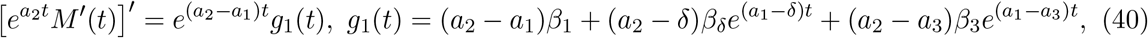

with *f* (0) = *M*” (0) > 0 and lim_*t*→∞_ *g*_1_(*t*) = (*a*_2_ − *a*_1_)*β*_1_ < 0. In terms of (39) and (40), we have (*a*_2_ − *δ*)(*a*_1_ − *δ*)*β_δ_* < 0 and (*a*_2_ − *a*_3_)(*a*_1_ − *a*_3_)*β*_3_ < 0, and so

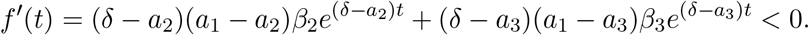 Hence, there exists a *t*_5_ > 0 such that *g*_1_(*t*) > 0 in (0, *t*_5_), *g*_1_(*t*_5_) = 0, and *g*_1_(*t*) < 0 in (*t*_5_, +∞). For *t* > 0 sufficiently large, *M*′ (*t*), dominated by the term with exp(−*a*_1_) in (32), is negative. We also recall that *M*′ (*t*) > 0 for *t* > 0 sufficiently small. Thus, there exists a finite *t*_6_ > 0 such that *M*′ (*t*) > 0 in (0, *t*_6_), and *M*′ (*t*_6_) = 0. Then *M*” (*t*_6_) ≤ 0, and (40) gives *f* (*t*_6_) = exp(*a*_1_*t*_6_)*M*” (*t*_6_) ≤ 0. It follows that *t*_6_ ≥ *t*_5_ and *f*(*t*) < 0 for all *t* > *t*_6_. Hence, by using (40) again, for each *t* > *t*_6_, we have

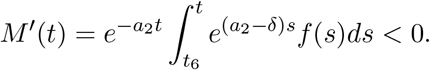
2. We present the proof for the case *a*_1_ < *a*_2_ < −*x*_2_ < *δ* < −*x*_1_ < *a*_3_, as the other cases can be handled similarly. With this specification, we have

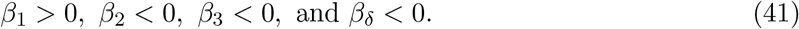 By (32) and (41), we obtain for all *t* > 0

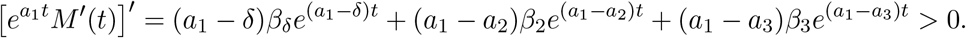 As *M*′ (0) = 0, it shows that 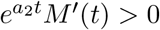, and so *M*′ (*t*) > 0 for all *t* > 0.
3. We give a short description for the proof with the additional condition that *x*_1_ < *x*_2_ < *δ* < *a*_1_ < *a*_2_ < *a*_3_, as the other cases can be proceeded in an analogous manner. Under this extra condition, (33) holds again. Thus the remaining discussion is the same as the proof given above for two complex valued *x*_1_ and *x*_2_, and we omit it here.

### Calculation of *M* (*t*) for cross-talking *n*-state model

Following the standard procedure of [15,33,39] we obtain the master equations for the cross-talking *n*-state model, which further leads to the differential system of gene state probabilities and the average mRNA level *M* (*t*) as follows

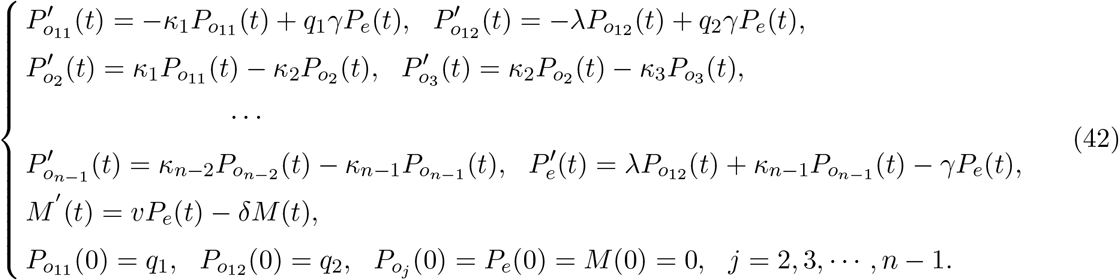

Here 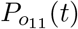 and 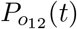 are probabilities of gene *off* 1 state selecting weak and signaling pathways, respectively. 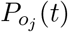, *j* = 2, 3, · · · , *n* − 1 are probabilities of gene *off j* states while the probability *P_e_*(*t*) is for gene *on* state. Also, we consider the initial condition for which the gene is totally *off* with no transcripts at the initial time.

For the special case of *q*_2_ = 0 which suggests that the gene activation is regulated by a single multi-step pathway, the system (42) reduces to

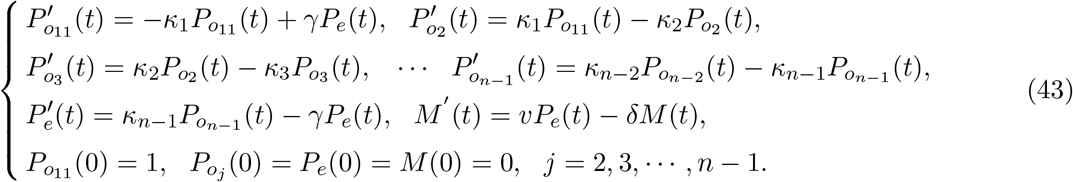

The crucial step to solve the system (42) or (43) is to calculate the eigenvalues of the coefficient matrix for equations of gene state probabilities [33, 39]. Here we illustrate a simple case of *κ*_1_ = *κ*_2_ = · · · = *κ_n_*_−1_ = *κ* in the system (43). Then its characteristic polynomial *f* (*x*) for gene state probabilities is expressed as

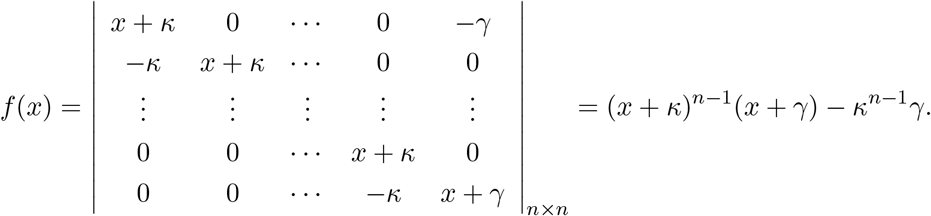

There is no standard method to solve the characteristic polynomial equation *f* (*x*) = 0 of degree *n* ≥ 5. However, the fundamental theorem of algebra shows that *f* (*x*) = 0 has *n* roots (eigenvalues) with complex roots always exhibiting in multiple pairs of conjugate complexes.

### The second moment of mRNA distribution for cross-talking three-state model (1)

By definition the second moment *μ*_2_(*t*) of the distribution of mRNA copy numbers is

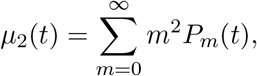

where *P_m_*(*t*) is the solution of master functions (2)-(6). To derive *μ*_2_(*t*), we need first calculate the partial mRNA average levels at each gene state, defined by

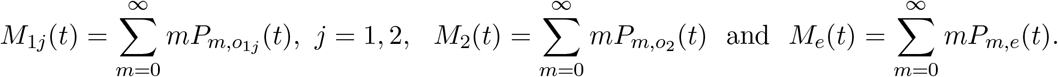

By these definitions and (2)-(6) we can derive the equations of *M*_11_(*t*), *M*_12_(*t*), *M*_2_(*t*), *M_e_*(*t*) and *μ*_2_(*t*) [33, 39]

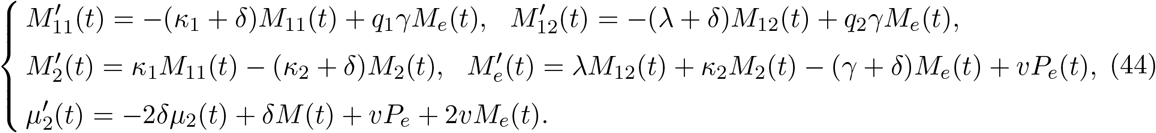

Here we focus on the initial condition

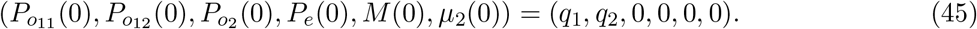

To solve the first-order differential system (44)-(45), we denote by

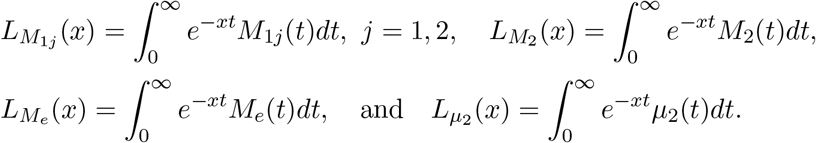

Then the application of Laplace transform to the first four equations of system (44) yields

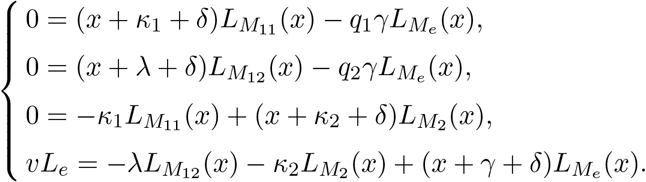

By applying Cramer’s rule in the theory of linear algebra to solve the above system, we obtain

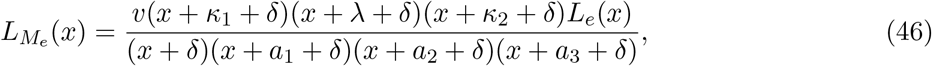

where *L_e_*(*x*) is given by (20) while *a*_1_, *a*_2_ and *a*_3_ are given by (15)-(19) or by the separate (22), (24) and (26) for the corresponding cases.

To solve the second moment *μ*_2_(*t*) from the system (44)-(45), we apply the Laplace transform to the last equation of (44), and then substitute (46) to obtain

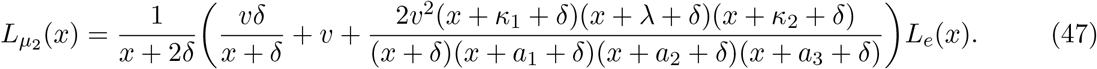

We note that *L_e_*(*x*) given by (20) can be simplified under the initial condition (12) or (45). The substitution of simplified *L_e_*(*x*) into (47) further gives

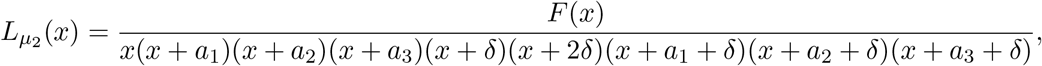

where *F* (*x*) takes the form of

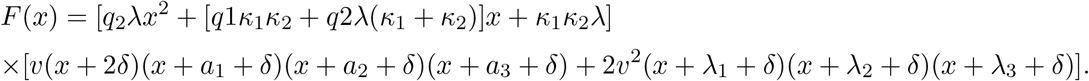

Under the condition 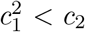 where *a*_1_, *a*_2_ and *a*_3_ are all positive real numbers, we decompose 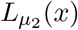 into the sum of partial fractions as

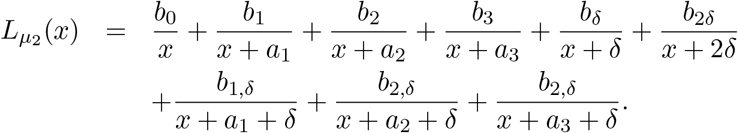

Then the application of the inverse Laplace transform to 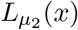 readily leads to

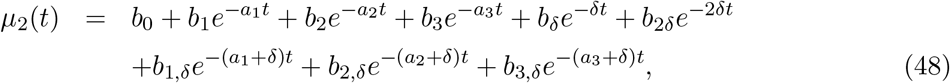

with *a*_1_, *a*_2_ and *a*_3_ being given by (22) and the coefficients taking forms of

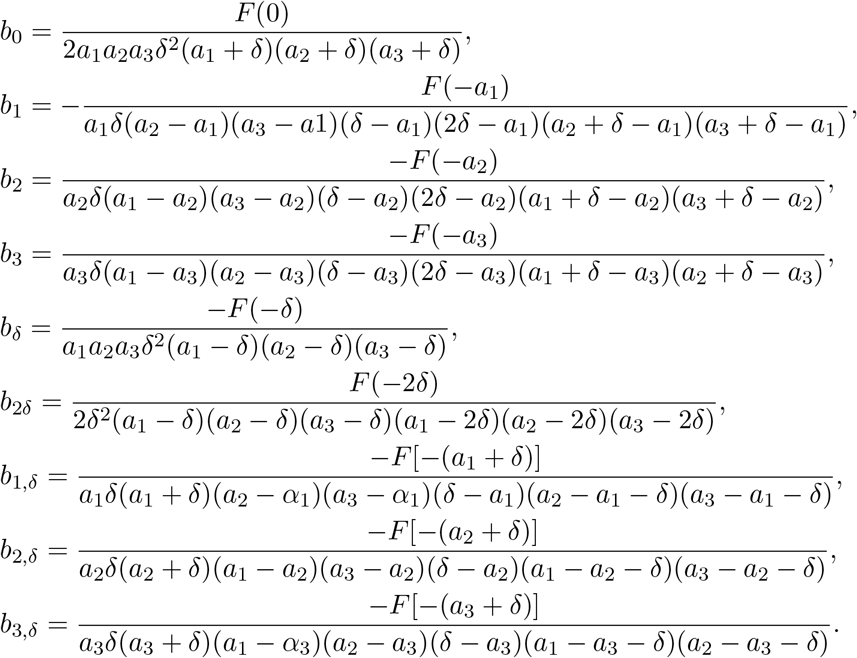

## Supporting Information

**Table S1.** Fitted system parameters for dynamical transcription data of mouse fibroblast genes.

**Fig. S1.** Oscillatory dynamics of mRNA average level *M* (*t*) generated by a single pathway with multiple sequential steps (*n*-state model). The increase in the step number *n* − 1 prolongs the initial response lag and enhances the following damped oscillation for *M* (*t*). Parameters are the fitted rates of *Cxcl1* gene in Table S1 with the average gene *off* duration *T_off_* being constant under (a) *κ_i_* = *iκ*_1_, *i* = 1, 2, · · · , *n* − 1; (b) *κ_i_* = *κ*_1_, *κ_j_* = 3*κ*_1_ for odd *i* and even *j*.

**Fig. S2.** Oscillatory dynamics of mRNA average level *M* (*t*) generated by the cross-talking *n*-state model. The increase in the step number *n* − 1 in the basal pathway has nearly no impact on the initial quick up-and-down dynamics of *M* (*t*) but significantly enhances the following gentle and damped oscillatory behavior. Parameters are the fitted rates of *Cxcl1* gene in Table S1 with the average gene *off* duration *T_off_* being constant under (a) *κ_i_* = *iκ*_1_, *i* = 1, 2, · · · , *n* − 1; (b) *κ_i_* = *κ*_1_, *κ_j_* = 3*κ*_1_ for odd *i* and even *j*.

## Author Contributions

L.C. G.L. and F.J. designed and performed the research; F.J. wrote the paper.

## Acknowledgements

This work is supported by Natural Science Foundation of China grants (Nos. 11871174; 12001127; 11631005), and by Program for Changjiang Scholars and Innovative Research Team in University (No. IRT 16R16). The funders had no role in study design, data collection and analysis, decision to publish, or preparation of the manuscript.

## Data Availability Statement

All relevant data are within the manuscript and its Supporting Information files.

